# *Vibrio* MARTX toxin processing and degradation of cellular Rab GTPases by the cytotoxic effector Makes Caterpillars Floppy

**DOI:** 10.1101/2023.04.19.537381

**Authors:** Alfa Herrera, Megan M. Packer, Monica Rosas-Lemus, George Minasov, John H. Brummel, Karla J. F. Satchell

## Abstract

*Vibrio vulnificus* causes life threatening infections dependent upon the effectors released from the Multifunctional-Autoprocessing Repeats-In-Toxin (MARTX) toxin. The Makes Caterpillars Floppy-like (MCF) cysteine protease effector is activated by host ADP ribosylation factors (ARFs), although the targets of processing activity were unknown. In this study we show MCF binds <u>Ra</u>s-related proteins in <u>b</u>rain (Rab) GTPases at the same interface occupied by ARFs and then cleaves and/or degrades 24 distinct members of the Rab GTPases family. The cleavage occurs in the C-terminal tails of Rabs. We determine the crystal structure of MCF as a swapped dimer revealing the open, activated state of MCF and then use structure prediction algorithms to show that structural composition, rather than sequence or localization, determine Rabs selected as MCF proteolytic targets. Once cleaved, Rabs become dispersed in cells to drive organelle damage and cell death to promote pathogenesis of these rapidly fatal infections.

## Introduction

Bacterial pathogens cause debilitating and lethal infections that are often difficult to treat. Understanding how to manage these infections is immensely important for improving quality of life and preventing death. The toxins secreted by extracellular bacteria are frequently the primary virulence factors required to establish colonization, promote maintenance, growth, and/or eventually dissemination. As such bacterial toxins are extremely important in pathogenesis and utilize several mechanisms of action to affect and manipulate host cellular processes to perform these roles.

*Vibrio vulnificus* produces a variety of secreted virulence factors to cause severe life-threatening gastrointestinal and wound infections, most commonly from the consumption of contaminated seafood or swimming with an exposed wound. The primary *V. vulnificus* virulence factor associated with sepsis and subsequent death is the composite Multifunctional-Autoprocessing Repeats-in-Toxin (MARTX) toxin encoded by the *rtxA1* gene ^1^. The MARTX toxin is a large polypeptide composed of conserved glycine-rich repeats at both the N- and C-termini between which there are variable effector domains and a Cysteine Protease Domain (CPD). The effectors and CPD are translocated across the eukaryotic host membrane through a pore formed by the repeats and released individually into the host cytoplasm. The MARTX toxin thus delivers multiple toxic virulence factors inside cells in a single bolus ^1,2^.

Across distinct *V. vulnificus* isolates, the effector content can vary from 2 to 5 effectors selected from a total of nine known effector domains ^3-5^. The effector domain most commonly present in *V. vulnificus* MARTX toxins is the Makes Caterpillars Floppy-like (MCF) domain. This 376 amino acid protein is found in most clinical isolates and is duplicated in some MARTX toxinotypes [Herrera, 2020 #904][Lee, 2019 #763][Herrera, 2020 #929]. Across the different toxinotypes, MCF is usually found directly after either the Actin Cross-linking Domain (ACD) or the Alpha/Beta Hydrolase (ABH) domain ^1^. Typically, MARTX effectors are released from the holotoxin by inositol hexakisphosphate (IP_6_) induced CPD processing. However, during autoprocessing, the N-terminus of MCF is not released due to lack of a peptidase recognition site. Instead, MCF remains tethered to the effector domain in front of it creating an “effector module” ^6^. MCF is an autoproteolytic cysteine protease and similar to other members of the C58 peptidase family, it has a Cys-His-Asp active site ^6-8^. Thus, instead of CPD-dependent release, MCF utilizes its autoproteolytic activity to release itself from the effector module and is required to fully process the MARTX toxin polypeptide into individual effector domains ^6^. Subsequent to autoprocessing, the exposed N-terminal glycine of MCF is acetylated ^8^. Combined, the autoprocessing and N-terminal acetylation results in the generation of “active” MCF (aMCF). aMCF induces the intrinsic apoptotic pathway, loss of mitochondrial membrane potential, Golgi destruction, and increases lethality in wound and gastrointestinal mouse models of infection ^6-9^.

MCF autoprocessing catalyzed within cells requires ADP ribosylation factors (ARFs) as a host stimulatory factor ^6,8^. ARFs cycle between active (GTP-bound) and inactive (GDP-bound) states to help regulate eukaryotic cell membrane trafficking ^10-12^. Depending on the isoform, they localize to specific subcellular locations including the Golgi, plasma membrane, and endosomes ^13-15^. There is a strong preference for induction of MCF autoprocessing by active ARFs, with ARF1 potentially the major activation factor within cells ^8^. However, ARFs are not the direct target leading to the cytotoxic effects as they are neither processed nor covalently modified following interaction with MCF ^6,8^.

In this study, we find MCF is a unique extracellular bacterial effector stimulated to activate by ARFs and subsequently act as a cysteine protease that directs the cleavage of <u>Ra</u>s-related proteins in <u>b</u>rain (Rab) GTPases to disrupt vesicular trafficking. The proteolytic activity of MCF is required to induce the degradation or cleavage of 50% of the Rabs tested independent of proteosome activation or caspase-3 mediated protein degradation. We resolve the structure of unbound aMCF in a conformational state that would allow for binding to Rabs directly. Furthermore, structure prediction analysis suggests MCF binds to Rabs and ARFs using the same interface. Recombinant protein assays confirm this and demonstrate MCF cannot simultaneously bind both GTPases and preferentially binds ARFs prior to autoprocessing. In addition, the C-terminal tail of Rabs is predicted to pass through a channel formed by aMCF containing its catalytic residues suggesting Rabs are cleaved at their tails, which is verified by mass spectrometry analysis. Ultimately, MCF mediated Rab degradation leads to mis-localization of endogenous Rab1B away from its expected perinuclear position.

## Results

### Rab4B is cleaved when co-expressed with MCF

Our previous work on MCF shows it is activated to autoprocess by host ARFs, but does not cleave or modify them to exert its cytopathic effects ^8^. Affinity purification mass spectrometry analysis on catalytically inactive MCF (MCF^CS^) by Lee *et al.* also demonstrated ARFs co-precipitate with MCF when ectopically expressed ^6^. Further examination of the published metadata by Lee *et al.* revealed significant detection of ten Rab proteins in the MCF interactome, albeit with a lower confidence score, than the ARF GTPases ^6^. In these data, Rab1B had the highest protein score (80.84) of the Rabs identified (Table 1). Since Rab GTPases share high structural similarity with ARFs and cells expressing MCF show extensive vesicular disruption ^8^, we postulated Rabs may be the biologically relevant cellular targets of the MCF effector domain.

**Table 1.** Catalytically inactive MCF (MCFCS)-interacting Ras-related proteins in brain (Rabs) identified by affinity purification mass spectrometry

|  |  |  |  |  |  |
| --- | --- | --- | --- | --- | --- |
| Protein | Molecular Weight (kDa) | Protein Score<sup>a</sup> | Coverage<sup>a</sup> | # of unique peptides<sup>a</sup> | Screen Results<sup>b</sup> |
| Rab1B | 22.2 | 80.84 | 46.77% | 3 | Degraded |
| Rab1A | 22.7 | 66.37 | 48.78% | 4 | Degraded |
| Rab5C | 23.5 | 60.7 | 71.76% | 7 | Cleaved |
| Rab2A | 23.5 | 54.72 | 47.64% | 8 | Unaffected |
| Rab7A | 23.5 | 47.19 | 62.32% | 9 | Unaffected |
| Rab11B | 20 | 44.68 | 54.75% | 8 | Cleaved |
| Rab21 | 24.3 | 34.99 | 37.33% | 7 | Cleaved |
| Rab10 | 22.5 | 31.25 | 23% | 3 | Unaffected |
| Rab14 | 23.9 | 27.22 | 38.6% | 6 | Degraded |
| Rab5B | 23.7 | 17.32 | 33.49% | 3 | Cleaved |

<sup>a</sup>Protein score, coverage, and number of unique peptides taken from data published by Lee et al. 6.
<sup>b</sup>Effect on Rab isoform based on data in Fig. 1b and Supplementary Fig. 1 and 2.

To initially test if Rabs are affected by MCF using reagents already in hand from prior studies ^16^, human embryonic kidney (HEK) 293T cells were co-transfected to express Green Fluorescent Protein (GFP)-Rab4B with MCF or catalytically inactive MCF (MCF^CA^). Although the expected size of full-length GFP-Rab4B is ∼50 kDa, the anti-GFP antibody in a western blot detected an additional smaller protein around 25 kDa only when co-expressed with catalytically active MCF (Fig. 1a). These data suggested at least Rab4B, and potentially other Rabs, could be the targets for MCF proteolytic activity.

**Figure 1.**
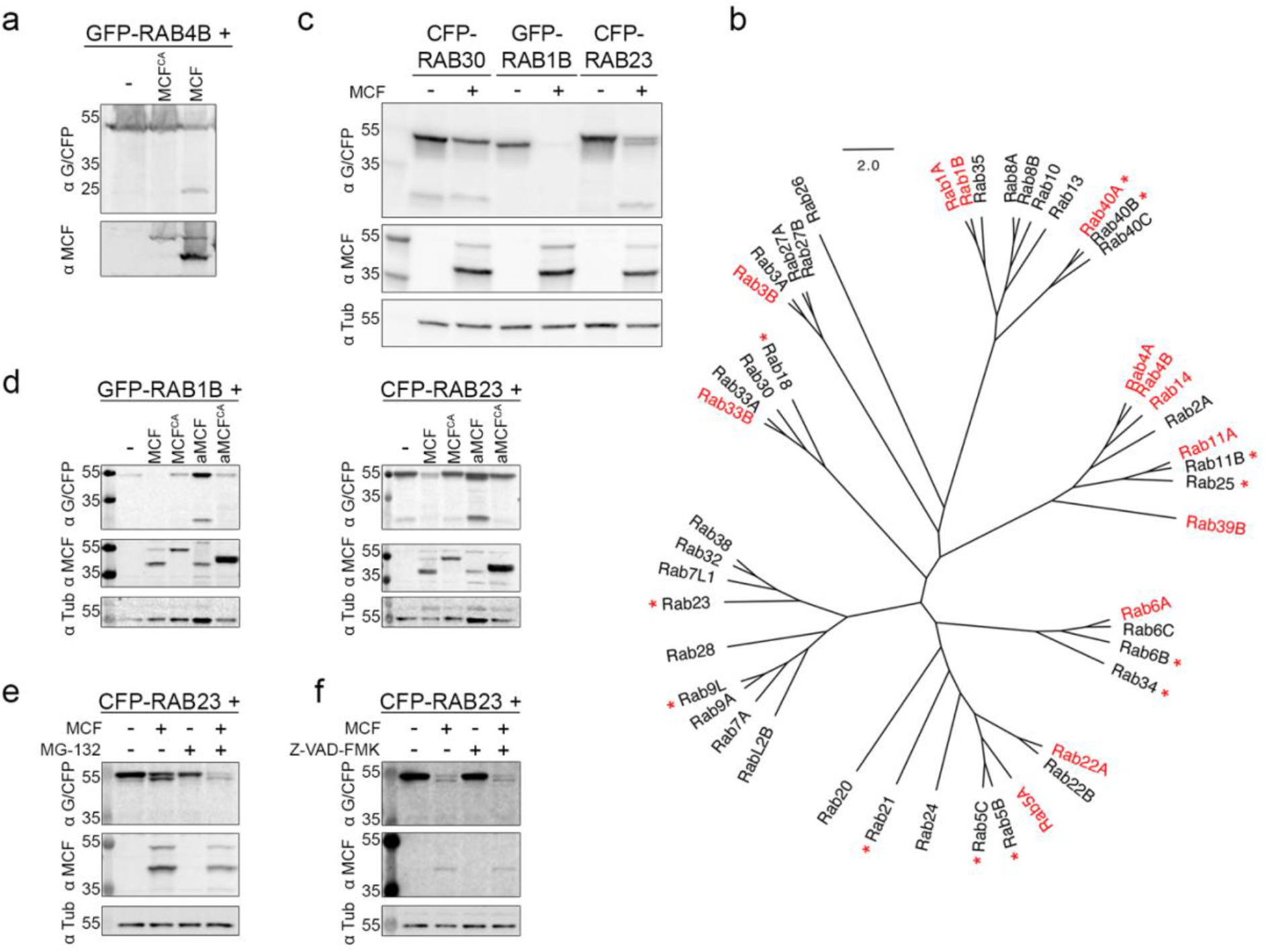
The cysteine protease activity of the MCF toxin induces variable degradation of Rab GTPases. **a, c** Fluorescently-tagged Rabs ectopically co-expressed in HEK 293T cells with MCF, MCFCA, or empty vector control (p3xFlag-CMV-7.1). Representative western blots shown of 35 μg of total protein from whole cell lysates recovered from cells (*n=3*). Rabs detected by anti-GFP/CFP (α G/CFP), MCF by anti-MCF (α MCF), and tubulin as a loading control by anti-α Tubulin (α Tub) antibodies. **b** Unrooted amino acid phylogenetic cladogram of Rab isoforms tested in the screen. Names in red indicate Rabs significantly degraded when co-expressed in HEK 293T cells with MCF, established by degradation factor determined by densitometry of bands on western blots from three independent experiments (see methods). Names with a red asterisk denote Rabs cleaved when co-expressed with MCF, as manually assessed. Western blots of independent transfections used to make the designations can be found in Supplementary Fig. 1 and analysis in Supplementary Fig. 2. **d** HEK 293T cells were co-transfected with MCF, MCFCA, aMCF, aMCFCA, or p3xFlag-CMV-7.1 and either GFP-Rab1B or CFP-Rab23 (*n=3*). Western blots on cell lysates recovered from these cells were performed as in (a). **e, f** Co-expression experiments and western blot analysis was completed as in (c) with CFP-Rab23 and MCF in the presence of either (e) 5 μM proteosome inhibitor MG-132 or (f) 5 μM pan-caspase inhibitor Z-VAD-FMK.

### MCF induces degradation and cleavage of at least 24 small Rab GTPase proteins

Nearly 70 unique Rab GTPases or splicing isoforms of Rabs have been identified in humans to date compared to only five ARF members ^17^. To assess if MCF was limited to Rab4B or more broadly interacts with Rabs, 48 fluorescently-tagged Rabs were co-expressed with MCF (Supplementary Fig. 1 and Supplementary Table 1). Percent degradation for each Rab co-transfected with MCF was quantified and compared to its parallel Rab co-transfected with an empty vector control (see methods). Rabs with a degradation factor 115% more than the standard deviation of its appropriate empty vector control were determined to be degraded as a result of MCF expression (Supplementary Fig. 2). This analysis revealed multiple distinct outcomes. In some cases, such as GFP-Rab1B, the GTPase was detectable only in the absence of MCF, indicating that the presence of MCF led to complete degradation of the GTPase (Fig. 1b names in red, 1c, and Supplementary Fig. 1 and 2). For other Rabs, such as with Cyan Fluorescent Protein (CFP)-Rab23, MCF co-transfection resulted in the appearance of a cleavage product slightly smaller than the expected size (Fig. 1b marked with red asterisk, 1c, and Supplementary Fig. 1 and 2). In some cases, such as Rab40A, the total amount of protein detected was reduced and a smaller band was detected, suggesting an intermediate where both cleavage and degradation are captured. Finally, not all Rabs, including CFP-Rab30, were affected by MCF co-expression as neither degradation nor a cleavage product were observed (Fig. 1b names in black, 1c, and Supplementary Fig. 1 and 2).

Overall, MCF induced significant degradation or cleavage of 24 of the 48 (50%) Rabs tested (Fig. 1b). Not surprisingly, some Rabs detected as potential targets were absent from the interactome reported by Lee *et al*. (Table 1) ^6^. In addition, not all Rabs identified in the interactome were degraded or cleaved in the co-expression screen suggesting MCF binding and cleavage may not be necessarily coupled ^6^.

### MCF selection of Rabs is independent of amino acid variation

We first speculated that MCF may select a subgroup of Rabs closely linked by amino acid similarity. This possibility was not supported since MCF affected Rab isoforms speckled across the phylogenetic tree (Fig. 1b). Notably, Rab2A is the only member in its nested clade unaffected by MCF co-expression (Fig. 1b and Supplementary Fig. 1). However, Rab2A is also the only construct in this clade cloned into the peCFP-C1 vector rather than peGFP-C2 (Supplementary Table 1). To confirm the nature of the tag did not alter Rab susceptibility to MCF, the coding sequence for Rab2A was re-cloned into peGFP-C2. Results verified Rab2A is not cleaved or degraded upon co-expression with MCF (Supplementary Fig. 1 and 2). Thus, overall amino acid sequence similarity between the affected Rabs did not confer the specificity of MCF.

### Rab GTPases are degraded during intoxication with MCF encoding *V. vulnificus*

To confirm the MCF induced Rab degradation observed was not an artifact of the overexpression system and occurs in the context of natural intoxication, HEK 293T cells were incubated at an multiplicity of infection of 5 for 2.5 hours with *V. vulnificus* strains that secrete a MARTX toxin with either no effectors or only the MCF effector domain ^9^. Proteins recovered from whole cell lysates were separated by SDS-PAGE and peptide sequencing was completed on an excised band encompassing all 20 - 30 kDa proteins (Supplementary Fig. 3a). Endogenous Rab5A, 5B, 7A, 11B, 14, and 21 all had fewer unique identified peptides in cells treated with bacteria that deliver MCF to cells compared to a toxin that delivers no effectors, including the complete absence of detected peptides for Rab 14, 21, and 11B when MCF was present (Supplementary Fig. 3b and 3c). Interestingly, there was not a significant difference in detected peptides during intoxication with the MCF encoding strain for Rab1A, despite it being degraded in the overexpression screen, while the opposite is true for Rab7A (Fig. 1b and Supplementary Fig. 3). Overall, these experiments confirm the capacity of MCF to cause Rab degradation when naturally delivered in a MARTX toxin by *V. vulnificus* with additional Rabs degraded when MCF is overexpressed in cells.

### MCF cysteine protease activity is required for Rab degradation

To determine whether the proteolytic activity of MCF is required for the Rab degradation and cleavage, we focused on Rab1B and Rab23 since (i) Rab1B is one of the most well studied Rab GTPases, is the top Rab interacting partner by Lee et *al*. ^6^, and is substantially degraded and (ii) Rab23 consistently generates a reproducible and discernable cleavage product when co-expressed with MCF. Furthermore, both play a role in autophagy, and an abundance of autolysosomes was previously observed in cells expressing MCF ^8,18-24^.

Cells were transfected with a plasmid construct that expresses protein to mimic MCF after autoprocessing (aMCF). Interestingly, degradation and cleavage were less efficient with aMCF co-expression than with MCF (Fig. 1d and Supplementary Fig. 4a). In addition, ectopic expression of catalytically inactive aMCF (aMCF^CA^) confirmed proteolytic activity of MCF is necessary (Fig. 1d and Supplementary Fig. 4a). The degradation and cleavage of GFP-Rab4B and CFP-Rab9L respectively was also confirmed to require the catalytic activity of MCF, and to occur most efficiently when transfected in a form prior to autoprocessing (Supplementary Fig. 4a). These data suggest a reorganization of MCF to aMCF proximal to Rabs is required for optimal targeting of Rabs.

These data do not exclude that MCF may indirectly activate the ubiquitin-proteosome pathway or apoptosis inside eukaryotic cells to induce the Rab degradation and cleavage. To address this possibility, co-transfection experiments were completed including the proteosome inhibitor, MG-132. In the presence of MG-132, CFP-Rab23 is cleaved when co-expressed with MCF, and thus is independent of proteosome activation (Fig. 1e and Supplementary Fig. 4b). Prior studies show MCF activates caspase-9, -7, and -3 and PARP-γ ^9^. To evaluate if MCF is causing Rab cleavage by stimulating caspase-3 mediated protein degradation, co-expression experiments were performed in the presence of a pan-caspase inhibitor. Inhibiting caspase activation did not prevent MCF induced CFP-Rab23 cleavage (Fig. 1f and Supplementary Fig. 4c). Combined these data support the hypothesis that the proteolytic activity of MCF is directly cleaving and degrading Rabs in host cells.

### MCF directly interacts with Rabs

To further support a direct cleavage model, we tested if MCF stably binds Rabs. Co-immunoprecipitation assays attempting to recover Rabs by precipitating MCF from cell lysates and vice versa from co-expression experiments were unsuccessful, suggesting their interaction is transient inside cells (Supplementary Fig. 4d). Moreover, catalytically inactive MCF^CA^ expressed in cells remains bound to ARFs, potentially hindering Rab binding and accounting for why Rabs were not detected in affinity purification mass spectrometry analysis by Herrera et *al*. and at low abundance by Lee et *al*. ^6,8^.

Therefore, an *in vitro* co-immunoprecipitation approach was used to test binding with recombinant glutathione *S*-transferase (GST)-tagged Rab1B or Rab23 and MCF. MCF and autoprocessed MCF (aMCF), consistently directly bound to either Rab. (Fig. 2a, 2b, and Supplementary Fig. 5a and 5b). In addition, the experiments show separately the C-terminal domain 1 of aMCF (aMCF^domain^ ^I^), consisting of residues 1-84, was not sufficient to bind to either Rab. Moreover, the N-terminal domain of aMCF (aMCF^domain^ ^II^)(residues 85-324), could also not bind to Rab1B or Rab23 alone. Additional assays completed confirm MCF and aMCF are capable of directly interacting with several Rab isoforms (Supplementary Fig. 6a - f). Interestingly, aMCF^domain^ ^II^ does bind to Rab5A and Rab11A (Supplementary Fig. 6e). Overall, these data suggest optimal binding to Rabs requires both the N- and C-termini of aMCF.

**Figure 2.**
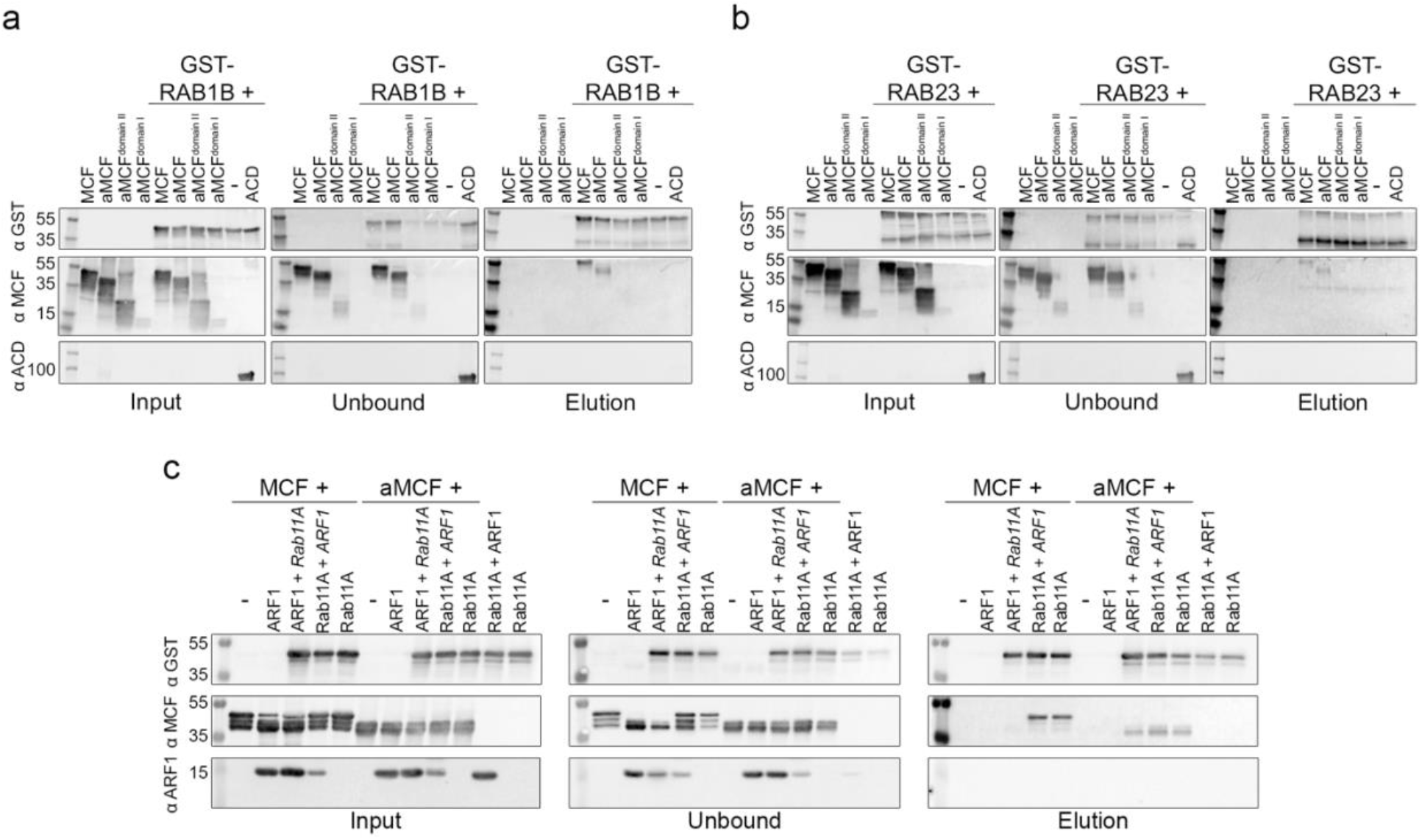
MCF and aMCF immunoprecipitate (IP) with Rabs. **a, b** Purified (**a**) GST-Rab1B or (**b**) GST-Rab23 were incubated with either MCF, aMCF, residues 85-324 of aMCF (aMCF^domain II^), or residues 1-84 of aMCF (aMCF^domain I^) at 37°C. Purified recombinant anti-ACD was used as a control for non-specific binding. Western blot on the incubated samples (input), flow through following incubation with GST beads (unbound), and sample following anti-GST IP (elution) using anti-MCF (α MCF), anti-GST (α GST), and anti-ACD (α ACD) antibodies. Representative gels (n=3). **c** Recombinant MCF or aMCF were incubated with either ARF1 or GST-Rab1B for two hours prior to the addition of the other GTPase followed by overnight incubation at 37°C. GTPase added after two hours in italics. Anti-GST IP and western blot on input, unbound, and elution samples were completed as above in (**a, b**) including anti-ARF1 (α ARF1) antibodies.

### MCF preferential binds ARFs prior to autoprocessing

Interestingly, none of the Rabs tested stimulated MCF autoprocessing *in vitro* (Fig. 2a, 2b, and Supplementary Fig. 6a). This is consistent with previous reports showing other Ras family GTPases, including ARF-like protein 4C (ARL4C) and secretion-associated Ras-related GTPase 1A (SAR1A), do not stimulate MCF autoprocessing ^6^. These data suggest ARFs and Rabs do not bind MCF simultaneously and thus ARF-stimulated autoprocessing of MCF occurs before Rab binding. To examine if ARFs and Rabs co-occupy MCF, co-immunoprecipitation assays were completed with ARF1 added either two hours before or after GST-Rab11A. Only when Rab11A was added prior to ARF1, did MCF co-immunoprecipitate with Rab11A, indicating a preference for MCF to bind ARF prior to autoprocessing (Fig. 2c). In contrast, aMCF co-immunoprecipitated with Rab11A regardless of whether ARF1 or Rab11A were added first, suggesting processed MCF does not prefer ARFs over Rabs (Fig. 2c). Of note, ARF1 is not simultaneously recovered in these reactions, showing MCF does not bind both GTPases simultaneously. These data indicate MCF interacts with ARFs and autoprocesses before binding to Rabs.

### Rabs are predicted to bind to the same interface of MCF as ARFs

To predict how aMCF binds to Rabs and subsequently causes their degradation or cleavage structure prediction software ColabFold was utilized ^25^. The resulting predicted aMCF model is nearly identical to the structure deposited for aMCF^CS^ (PDB code 6ii6) ([RMSD] = 0.534 Å for 348 Cα atoms) in complex with ARF3^Q71L^(Supplementary Fig. 7a) ^6^. The predicted aMCF is also composed of a helix bundle domain at its N-terminus and an α/β -fold domain at its C-terminus^6^. The predicted Rab1B structure also aligns well with the solved structure of Rab1B^4-175^ (PDB code 4i1o) ([RMSD] = 0.582 Å for 172 Cα atoms) (Supplementary Fig. 7b) ^26^. The analysis predicts Rab1B interacts in the same manner as ARF3^Q71L^ with closed aMCF (Fig. 3a and Supplementary Fig. 7c). Most of the interaction is predicted to occur between the interswitch and switch 2 regions of Rab1B and the α4 and α7 helices of aMCF (Supplementary Fig. 7d). Just as with ARF3^Q71L^, when bound to Rab1B, the N-terminal α-helices of aMCF fold toward the α/β -fold domain, and the site containing the catalytic cysteine is exposed. The analysis predicting both GTPases bind to the same interface of aMCF supports the co-immunoprecipitation experiments demonstrating they competitively inhibit the binding of each other.

**Figure 3.**
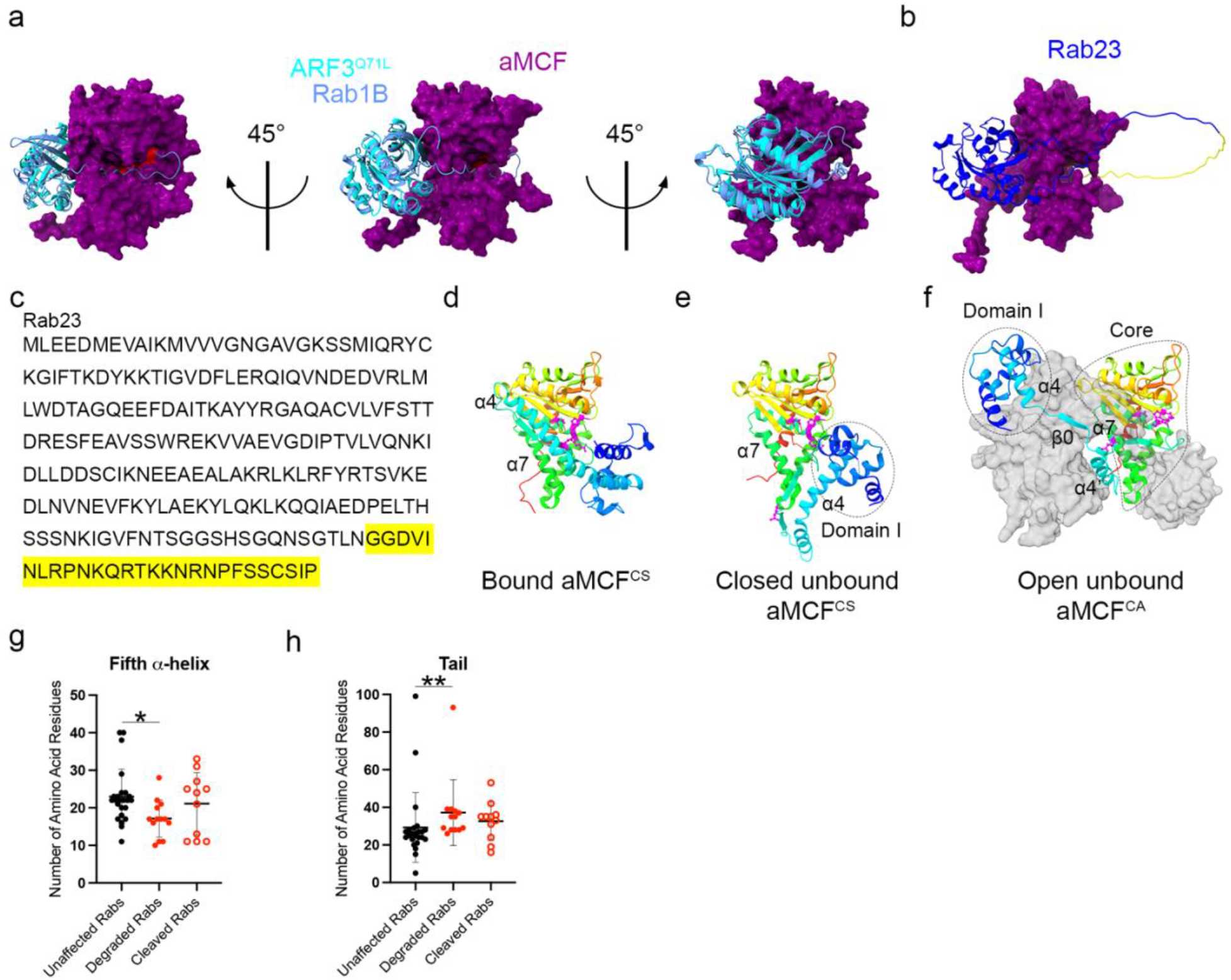
Structure of unbound aMCF toxin and predicted complex structure of aMCF with Rabs. **a** Three views of predicted complex ribbon structure of Rab1B (cornflower blue) bound to surface representation of aMCF (purple) overlayed onto the ribbon structure of ADP Ribosylation Factor 3 mutated to mimic the active state (ARF3^Q71L^) (cyan) bound to aMCF^CS^ (not shown) (PDB code 6ii6) ^6^. Residues important for proteolytic activity of aMCF (Gln118, Arg132, Cys133, Asp134, His245, and Asp264) highlighted in red. **b** Projected ribbon structure of Rab23 (blue) in complex with surface representation of aMCF (purple). Residues removed during co-transfection with MCF in HEK 293T cells are highlighted in yellow. **c** Protein sequence of Rab23 designating residues cleaved off (shaded in yellow) during co-expression with MCF in HEK 293T cells. **d**, **e, f** Ribbon structure of (**d**) closed aMCF^CS^ derived from the complex structure with ADP Ribosylation Factor 3 mutated to mimic the active state (ARF3^Q71L^) (PDB code 6ii6) ^6^, (**e**) closed unbound aMCF^CS^ (PDB code 6ii0) ^6^, and (**f**) open unbound aMCF^CA^ (PDB code 8SFG) with surface representation of swapped dimer in light gray, in a rainbow spectrum from the N-terminus in blue to the C-terminus in red. α4 and α7 helices (according to PDB code 6ii6) ^6^ important for flexibility and binding denoted and distinct domains of unbound aMCF outlined. Residues important for proteolytic activity of aMCF (Gln118, Arg132, Cys133, Asp134, His245, and Asp264) are highlighted in magenta on structures ^6-8^. **g, h** Length in number of amino acid residues of the (**g**) fifth α- helix and (**h**) hypervariable tail in Rabs screened determined from structure prediction models. Significance was calculated using a non-parametric Kruskal-Wallis test, P value: 0.1234 (ns), 0.0332 (*), 0.0021 (**), 0.0002 (***), < 0.0001 (****).

### aMCF forms a channel through which the hypervariable C-terminal tail of Rab1B and Rab23 inserts

A surprise of the ColabFold analysis was that the best model predicted 14 residues of the flexible C-terminal hypervariable region domain, or tail, of Rab1B inserts into a channel formed by aMCF^domain^ ^II^ and aMCF^domain^ ^I^ (Fig. 3a and Supplementary Fig. 7d). Notably, at the surface this channel contains the catalytic residues Arg132, Cys133, Asp134, and His245 that are essential for the proteolytic activity of aMCF, as well as Gln118 which may function as an oxyanion residue. However, the catalytic residue Asp264 was not modelled to be at the surface of aMCF in the channel (Fig. 3a and Supplementary Fig. 7e). In addition, ColabFold predicted aMCF binds to Rab23 similarly to Rab1B, and suggests its tail also passes through the channel formed by aMCF (Fig. 3b). The structure prediction analysis therefore suggests aMCF is directly cleaving Rabs at their C-terminal tails, potentially disrupting activity by removing regions essential for membrane binding, localization, and function of Rabs ^27-30^

### CFP-Rab23 is cleaved at its C-terminus when co-expressed with MCF

To identify the exact cleavage site on Rabs within cells we concentrated on CFP-Rab23 which consistently showed a detectable cleavage product resulting from co-expression with MCF (Fig. 1c and Supplementary Fig. 1) and was detected with its N-terminal CFP-tag. Affinity purification mass spectrometry analysis conducted on CFP-Rab23 recovered from cells co-transfected with MCF (Supplementary Fig. 8a) revealed cleaved CFP-Rab23 had less coverage at the C-terminus than the uncleaved bands (Supplementary Fig. 8b). Peptide analysis further showed CFP-Rab23 is cleaved between Asn210 and Gly211, removing the last 27 amino acids at the C-terminus of the protein critical for its proper functioning (Fig. 3b, 3c (residues in yellow), and Supplementary Fig. 8b and 8c).

While CFP-Rab23 is cleaved when co-transfected with MCF, not all is truncated as protein corresponding to the full-length size is still detectable (Fig. 1c). Rabs are typically post-translationally modified at their C-terminal cysteines and may also be modified at other internal residues ^31-34^. In fact, two-dimensional gel analysis showed CFP-Rab23 separates into five discrete populations with different isoelectric points, representing five distinct populations of post-translationally modified Rab23 (Supplementary Fig. 8d, left). However, when co-expressed with MCF, only three distinct populations of full-length CFP-Rab23 are detectable (Supplementary Fig. 8d, right). Furthermore, coincident with a shift in electrophoretic mobility for cleaved CFP-Rab23, two new populations with different isoelectric points are detectable. These data indicate only two forms of post-translationally modified CFP-Rab23 are cleaved with MCF co-expression.

### aMCF transitions from a closed to an open conformation following activation

While the structure prediction software was highly informative as it predicted Rabs to be cleaved at their C-termini by passing through the channel formed by MCF, there are inconsistencies with this model at the catalytic site. A key proteolytic residue, Asp264, is not exposed in the channel in this conformation of aMCF and does not align with a scissile bond in Rab1B (Fig. 3a, 3b, and Supplementary Fig. 7e and 10a). The predicted organization of the catalytic residues of aMCF bound to Rab1B, which mimic it bound to ARF3^Q71L^, would not allow for aMCF to function and cleave (Fig. 3a, 3d, and Supplementary Fig. 7c and 10a). The previously solved structure of unbound aMCF^CS^ (PDB code 6ii0) ^6^ also shows a channel, although more tightly closed and narrow (Fig. 3e and Supplementary Fig. 9a and 9b). In this conformation of aMCF, the Gln118 residue is located away from the channel and the catalytic residue Asp264, while slightly more exposed on the surface of the protein, is oriented behind the other catalytic residues. (Fig. 3e and Supplementary Fig. 9a and 9b, residue in yellow). This argues MCF may sample an alternative conformation in which the channel is more open and catalytic residues more exposed.

To assess alternatives for a structure of unbound aMCF in the absence of ARF or Rab, the crystal structure of unbound aMCF^CA^ (PDB code 8SFG) at 2.8 Å was determined using purified recombinant protein labeled with selenomethionine (SeMet) for structure determination by single-wavelength anomalous diffraction (SAD). The conformation of aMCF^CA^ obtained was a swapped dimer (Fig. 3f) and differed from the previously published unbound aMCF^CS^ (PDB code 6ii0) ^6^ (Fig. 3e) or the structure of aMCF^CS^ in complex with ARF3^Q71L^ (PDB code 6ii6) ^6^ (Fig. 3d). While the three aMCF structures differ in their states, their overall composition is similar (PDB code 8SFG to PDB code 6ii6: [RMSD] = 0.711 Å for 334 Cα atoms; PDB code 8SFG to PDB code 6ii0: [RMSD] = 0.76 Å for 326 Cα atoms) (Supplementary Fig. 9c and 9d).

This structure suggests aMCF monomers adopt an “open” conformation consisting of the N-terminus reorganized away from the body of MCF, and tethered to the C-terminal domain by an unstructured region (Fig. 3f). This unstructured, linker region (residues 83-118) is created in part by “unwinding” of α4 (according to closed bound aMCF, PDB code 6ii6) ^6^, into a new β0 strand surrounded by two shorter α4 and α4’ helices (Fig. 3a and 3f). At the N-terminus the first 82 residues of open aMCF^CA^ (aMCF^domain^ ^I^) consist of a four-helical bundle (α helices 1 - 4) with one charged end. At the C-terminus, the larger “core” domain of open aMCF^CA^ (aMCF^core^), following the linker, consists of 243 amino acids and contains the catalytic residues (Fig. 3f). Most of the proteolytic residues of open aMCF^CA^ are at the surface of the protein and very exposed (Fig. 3f and Supplementary Fig. 9a and 9e). Surprisingly, the catalytic Asp264 residue remains buried even in this conformation. The open aMCF^CA^ structure solved corresponds to aMCF following autoprocessing and release from ARFs, in an altered conformational state. Combined the structures suggest the α4 helix of aMCF is highly flexible and allows unbound aMCF to transition from a closed to an open conformation. We suggest the open conformation of aMCF is more favorable for binding Rabs.

### Rab C-terminal structure and flexibility correlates with susceptibility for MCF induced degradation

According to the screen, specificity of affected Rabs is not based on amino acid homology (Fig. 1b). Furthermore, affected Rabs do not localize to any particular subcellular region within the cell and are found ubiquitously throughout ^17^. We considered whether the affinity of MCF to select Rabs may account for their susceptibility to cleavage or degradation. Considering structure prediction analysis indicated most of the binding between the proteins occurs at the G domain of Rabs, we examined whether residue differences in this region correlated with susceptibility. aMCF-Rab complex structures generated in AlphaFold2 overlayed onto the aMCF^CS^-ARF3^Q71L^ complex indicated the 22 residues identified to be essential in the G domain on ARF3^Q71L^ for binding to aMCF tend to be functionally conserved at those positions in the Rabs from the screen (Supplementary Fig. 10) ^6^. Furthermore, there are no significant differences at these 22 residues between the degraded and cleaved Rabs and those that are not affected. This analysis supports the *in vitro* co-immunoprecipitation data showing MCF is capable of binding to several Rab isoforms (Fig. 2, and Supplementary Fig. 5, 6, and 7). Of note, the predicted complex structure of aMCF with one degraded (Rab40A), one cleaved (Rab40B), and six unaffected Rabs (Rab L2B, Rab40C, Rab2A, Rab6C, Rab13, and Rab20) did not align well with the complex of ARF3^Q71L^-aMCF^CS^ ([RMSD] = 0.657 - 0.836 Å for 246 - 288 Cα atoms) and were not included in the analysis (Supplementary Fig. 10). Overall the predicted co-structures of MCF with Rab isoforms in the screen indicate residue differences in the G domain do not account for their susceptibility.

Alternatively, since the structure prediction analysis suggested MCF cleaves Rabs at their C-termini and this is confirmed for CFP-Rab23, Rab tails were examined more closely in the predicted models for structure and length associated to specificity. The mean of the number of residues that make up the fifth α-helix from which the C-terminal tail extends is 17.15, 21.09, and 22.95 amino acids for degraded, cleaved, and unaffected Rabs respectively (Fig. 3g, Supplementary Fig. 11, and Supplementary Table 1). Thus, the average length, in amino acid residues, of the α5 helix is significantly shorter in degraded Rabs compared to those unaffected. Furthermore, while not statistically significant, this helix tends to be bent to a larger degree in unaffected Rabs (7.7°) than in degraded (4°) or cleaved (4.5°) Rabs (Supplementary Table 1). AlphaFold2 predicts a short break disrupts this helix in four of the 24 affected Rabs (∼17%) however, 11 of the 24 unaffected Rabs (46%) have a break in the helix. The C-terminal tail of degraded Rabs (mean 37.23) is made up significantly more residues than the tail of unaffected Rabs (mean 29.21) (Fig. 3h, Supplementary Fig. 11, and Supplementary Table 1). There is predicted to be secondary structure in 46.2% (6/13) degraded, 18.2% (2/11) cleaved, and 33.3% (8/24) of unaffected Rabs. Altogether, these data suggest both the length, flexibility, and structure of Rabs at their C-terminus is associated with their susceptibility to be degraded or cleaved when co-expressed with MCF.

### Expression of MCF disrupts Rab1B localization

Modifications to the C-termini of Rabs alter their subcellular localization ^27,28,30^. Furthermore, siRNA knockdowns of Rab1B result in GBF1 dispersal and its loss in co-localization with trans-Golgi marker GM130 ^35^. Importantly, these knockdowns also result in dissolution of the Golgi, similar to that observed in cells ectopically expressing MCF in prior immunofluorescence and transmission electron microscopy studies ^6,8^. To examine if MCF alters endogenous Rab1B localization, immunofluorescence microscopy was performed to observe the GTPase in host cells. Endogenous Rab1B in Cos7 cells transfected with an empty vector control, localized tightly at its expected perinuclear location (Fig. 4 and Supplementary Fig. 12). However, endogenous Rab1B in cells expressing MCF-eGFP, was no longer condensed or positioned next to the nuclei. MCF expression in Cos7 cells degrades Rab1B, dispersing it throughout the cell.

**Figure 4.**
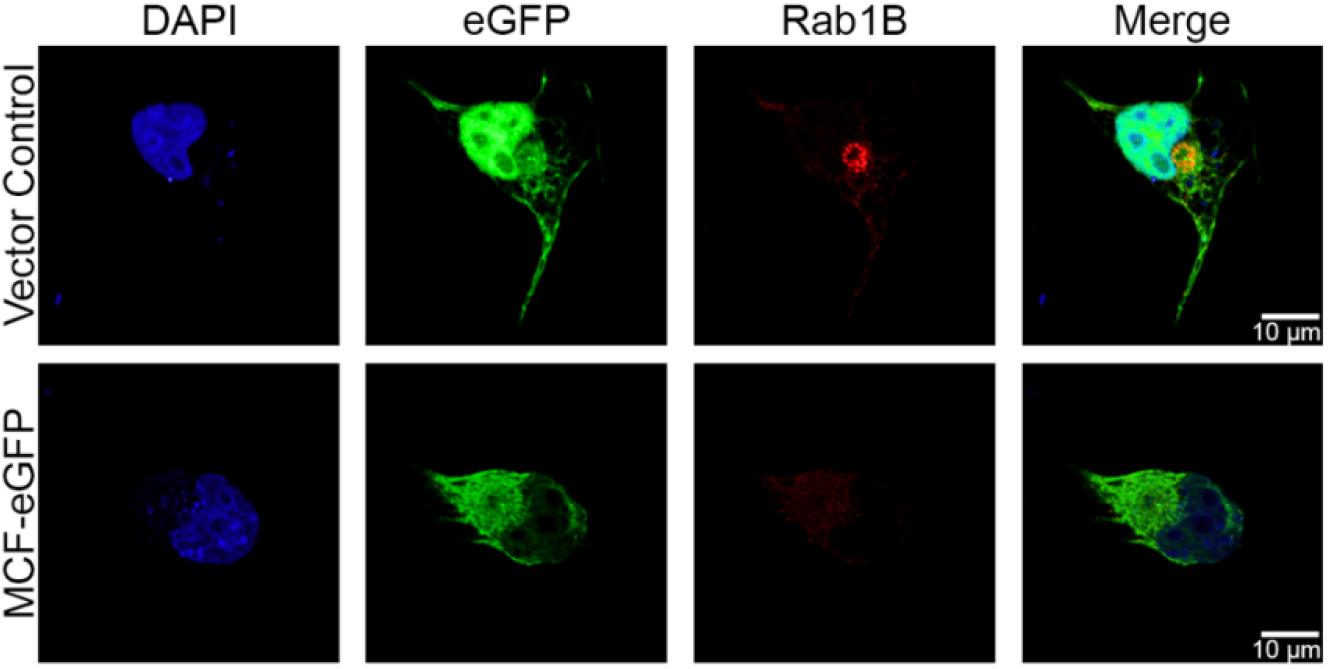
Ectopic expression of MCF mis-localizes endogenous Rab1B. Immunofluorescence microscopy of Cos7 cells transfected with empty vector control (peGFP-N3) or MCF-eGFP (green) for 18 hours, fixed, and stained for 4′,6′- diamidino-2-phenylindole (DAPI) (blue), and endogenous Rab1B (red).

## Discussion

The number of bacterial toxins discovered to specifically direct their activity toward altering Rab biology is on the rise. SidM (also known as DrrA), LepB, and Lgp0393 of *Legionella pneumophila*, SopE, SopE2, and SopD_2_ of *Salmonella enterica*, EspG of Enteropathogenic *Escherichia coli*, and VirA of *Shigella flexneri* mimic host regulators that alter the Rab activation state ^36-49^. Other effectors covalently modify Rabs through AMPylation (*L. pneumophila* SidM), de-AMPylation (*L. pneumophila* SidD), phosphocholination (*L. pneumophila* AnkX), dephosphocholination (*L. pneumophila* Lem3), glucosylation (*L. pneumophila* SetA), N-acetylglucosaminylation (*Yersinia mollaretii* YGT and *S. enterica* SseK_3_), ADP-ribosylation (*Y. mollaretii* YART and *Pseudomonas aeruginosa* ExoS), and ubiquitination (*L. pneumophila* SidE, SdeA, SdeB, and SdeC). All of these covalent modification prevent nucleotide exchange or interaction with GAPs, GEFs, or Rab effectors ^39,44,45,47,50-60^. However, only the Type III secretion effector GtgE of *S. enterica* has been found to directly cleave Rabs to date ^48,61-63^. The protease GtgE targets only a handful of nearly identical Rabs (Rab32, 38, 29).

The modifications on Rabs by these effectors occur mostly at the G domain, commonly the switch II domain, and target only a small subset of Rabs. Thus far, these effectors are predominantly encoded by intracellular pathogens to help form bacterium containing vacuoles, modulate vesicle trafficking, block host protein secretion, and inhibit phagolysosome maturation to increase their survival within the host and facilitate infection.

In this study, we show unlike any other effectors that target Rabs, MCF is delivered from a toxin expressed by an extracellular bacterium to cleave and degrade a broad range of Rab GTPases, which ultimately results in fragmentation of the Golgi and mitochondrial damage (Fig. 5). Co-immunoprecipitation assays show MCF directly binds to Rabs and this is supported by structure prediction analysis. Furthermore, considering the proteolytic activity of MCF is required for Rab degradation and cleavage in cells while proteosome and caspase activation are not, these data suggest MCF is directly cleaving Rabs. However, *in vitro* assays completed thus far using recombinant proteins have not shown direct cleavage of Rabs by MCF. In cells, Rab tails are post-translationally modified which facilitates insertion of the tail into lipid bilayers to anchor G domains to membranes ^34,64^. Furthermore, modified Rab tails are hydrophobic and are typically masked while cytosolic through interactions with GDP dissociation inhibitors (GDIs) until membrane insertion ^65,66^. The lack of post-translational modification(s) on Rabs and/or membrane binding, in our recombinant protein assays may account for the inability to detect direct cleavage by MCF thus far. In fact, two-dimensional analysis indicates only specific populations of post-translationally modified Rab23 are targeted for degradation by MCF. Alternatively, an additional unknown host factor may be required for Rab cleavage to occur.

**Figure 5.**
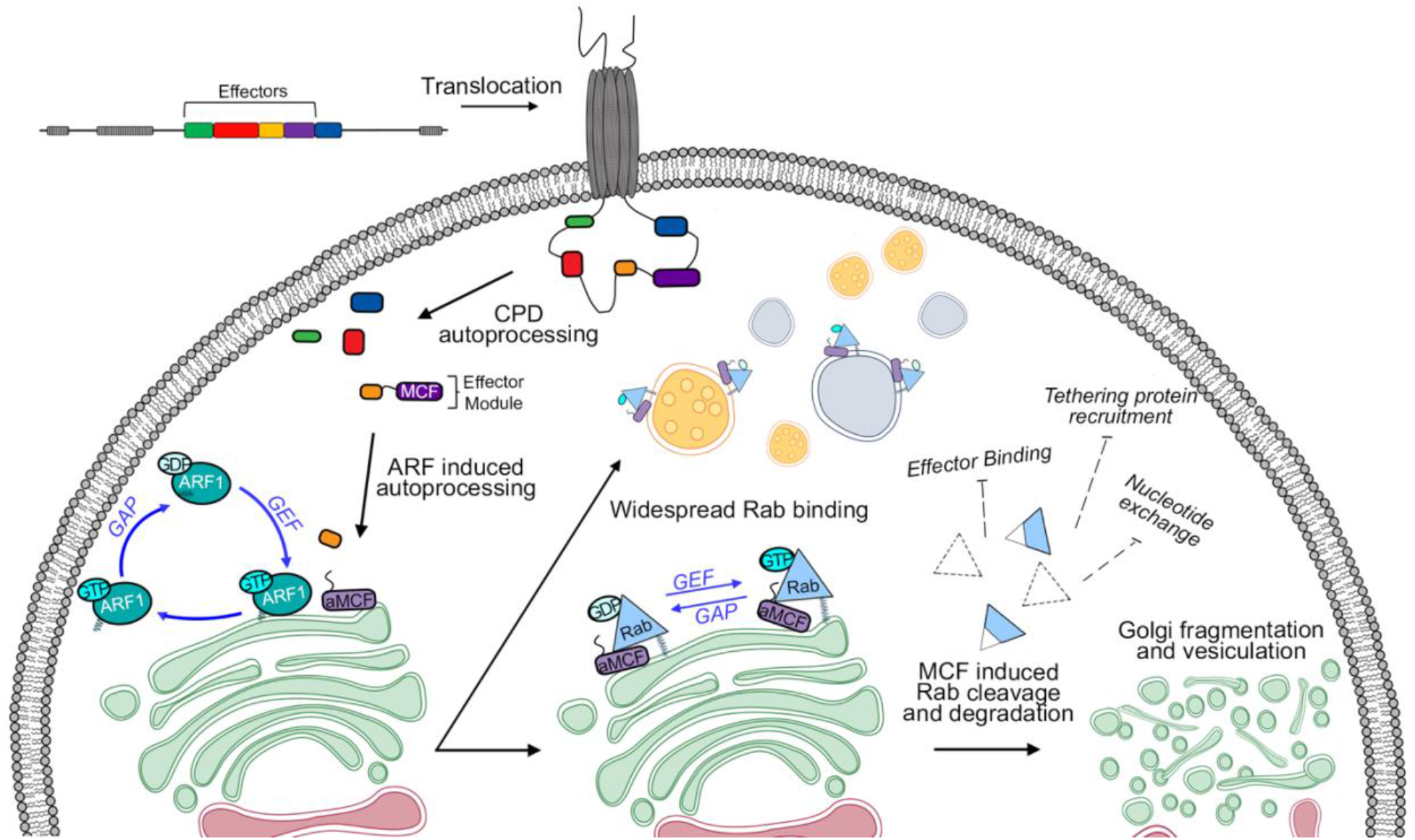
Model of MCF mediated Golgi fragmentation. After translocation, cysteine protease domain (CPD) processing releases individual effectors and MCF effector module. Binding to GTP-bound ARFs (ARF1 shown) stimulates MCF autoprocessing to release aMCF. Following release aMCF binds phosphatidylinositol-5-phosphate and is N-terminally acetylated. aMCF binds many membrane bound Rabs, with their hypervariable tail exposed (undetermined whether dependent on the nucleotide bound status of Rabs). aMCF induces widespread degradation and cleavage of Rabs which could prevent proper functioning. Rab dysfunction results in Golgi fragmentation. Golgi and ER modified from BioRender.com.

There are disparities in Rabs capable of interacting with MCF and Rabs whose degradation is induced by MCF. Furthermore, since there is no difference between unaffected and degraded Rabs at key binding residues, our data predicts Rab binding is not coupled to degradation. A key of our study was discovery that 50% of Rabs tested were susceptible to MCF under the conditions tested. It is notable that our analysis could have both over- and/or under-estimated the number of Rabs susceptible to MCF since the population of Rabs degraded may depend on the experimental conditions and cell type used. The ectopic expression system used produces substantially higher levels of proteins and thus may increase localization at atypical subcellular locations than would be found endogenously in HEK 293T cells with naturally delivered MCF leading to an over-representation of Rabs. Alternatively, some Rabs may not have been properly localized in our experimental conditions and thus under or over represented. Additional biophysical and cell biological studies are needed across the expanse of Rabs to define the relevant Rabs degraded during natural models of wound and gastrointestinal infections.

The sequential binding to GTPases by MCF suggested by the *in vitro* co-immunoprecipitation assays and prior studies establishing MCF releases ARFs following activation in cells, align with a hypothesis of a tightly regulated activation of MCF ^8^. Still, how the structural conformation of MCF transitions in binding from ARFs to Rabs is not clear. Our solved structure of unbound aMCF available for Rab binding provides new insights on the flexibility and reorganization of aMCF following the release of ARFs. However, the structure prediction analysis indicating Rabs bind to closed aMCF conflicts with the anticipated binding of open aMCF with Rabs. Therefore, we propose two potential models in which Rabs either bind to closed aMCF or open aMCF. It is possible our solved open aMCF structure is an artifact of crystallization and thus, Rab1B via its interswitch and switch 2 regions binds to closed aMCF at its α4 and α7 helices as predicted by the computational models (Fig. 6, Model 1). Alternatively, if our open aMCF structure is correct, alignment of open aMCF with the predicted Rab1B-MCF structure ([RMSD] = 0.667 Å for 209 Cα atoms) suggests Rab1B, using mostly its switch 1 and 2 regions, binds to the α7 helix of open aMCF (Fig. 6, Model 2A). However, this model does not account for the presence of the linker region formed from the rearrangement of the α4 helix of closed aMCF into a small β sheet between the α4 and α4’ helices of open aMCF, which would obstruct Rab binding in this manner. Therefore, we used Alphafold2 to predict how Rab1B would bind to open aMCF^core^, residues 119 – 361 (Supplementary Fig. 13). Alignment of open aMCF with the predicted Rab1B-aMCF^core^ structure ([RMSD] = 0.779 Å for 208 Cα atoms) suggests binding occurs similarly with the α7 helix of open aMCF. Rab1B is still predicted to bind mainly via its switch 1 and 2 regions, but is tilted slightly upward and forward (Fig. 6, Model 2B). In the second model, following Rab binding, open aMCF may then use its linker region to swing the aMCF^domain^ ^I^ over the tail of Rab1B and clamp it into the proteolytic channel or remain open.

**Figure 6.**
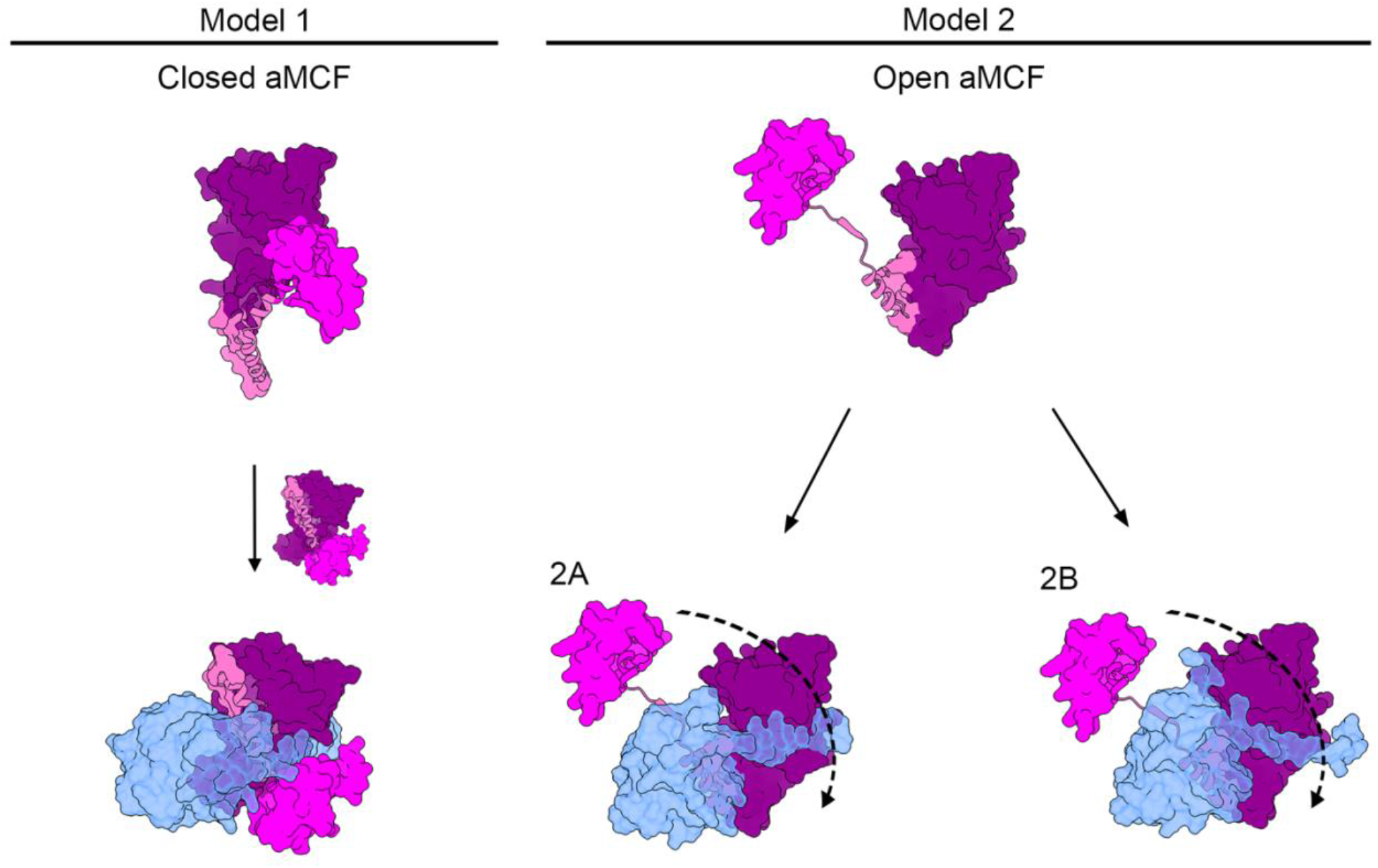
Proposed models of Rab1B binding based on solved structures of open and closed autoprocessed MCF. N-terminal domain I (residues 1 - 82), linker (residues 83 - 118), and the core domain (residues 119 - 361) of open aMCF in magenta, hot pink, and purple respectively and Rab1B in cornflower blue. Models shown as surface structures with the exception of α4 of closed aMCF as it transitions into α4, β0, and α4’ of open aMCF presented as a ribbon structure. In model 1, closed unbound aMCF (PDB code 6ii0) ^6^ transitions to a structure that resembles its ARF3-bound conformation (PDB code 6ii6) ^6^ to bind Rab1B, as is predicted in Fig. 3a. In model 2, Rab1B binds to open aMCF (PDB code 8SFG), as overlayed onto the (2a) predicted Rab1B-aMCF (Fig. 3a) or the (2b) predicted Rab1B-aMCF^119-361^ co-structure (Supplementary Fig. 13).aMCF^domain^ ^I^ may then swing over the tail of Rab1B and clamp it into the proteolytic channel or remain open.

Considering both Arg132-Cys133-Asp134 and Cys133-His245-Asp264 have been proposed as the catalytic triad of MCF ^6,7^, it is interesting residue Asp264 is mostly occluded from the surface of aMCF in the predicted and solved structures (Fig. 3a, 3b, 3d-f, and Supplementary Fig. 7e, 10a, 10b, and 10e). Asp264 and His245 have been shown to be important for MCF autoprocessing *in vitro*, but are not essential for its ability to induce cell rounding. The RCD residues are mostly exposed and found in the channel of aMCF, in addition to being important for autoprocessing and its cytopathic effects, strengthening the hypothesis these are the key residues for MCF catalysis.

Despite their similarity, Rabs localize to discrete regions inside the cell, are not usually functionally redundant, nor interact with the same effectors ^65,67,68^. Data indicating the fifth α- helix and the hyper variable domain tail of Rabs are important for their specificity and localization is emerging ^30,68,69^. Gyurkovska *et al.* recently showed despite Rab1A and Rab1B sharing 92% amino acid similarity, only Rab1A is required for stress-induced autophagy ^29^. Moreover, replacing the tail of Rab1B with that of Rab1A enables Rab1B to regulate autophagy. This corresponds with structural analysis of the C-termini of Rabs from the predicted structures suggesting the length, flexibility, and conformation at this end contributes to their susceptibility for MCF-induced degradation. MCF preferentially directs the cleavage of Rabs with a fifth α- helix and tail approximately 17 and 37 amino acid residues in length respectively. The fifth α-helix of a Rab cannot be too long as this decreases the proximity of the tail to the proteolytic channel formed by aMCF, which can be further impaired by a short tail which would be unable to cross through. An example is Rab35, an unaffected outlier in its clade, which has an α5 helix 24 residues longer and its tail ∼24 residues shorter than the other members in its clade. Rab6C has a tail almost twice the length of the others in its clade, which are affected in the screen, possibly making it more difficult for aMCF to correctly orient the tail for cleavage. However, helix and tail length alone do not account for susceptibility. Rab2A has a similar tail and helix length within its clade but is unaffected. However, Rab2A does have a different arrangement at its C-terminal cysteines with an “XXXCC” instead of a “XXCXC” motif. These data indicate Rab susceptibly could be complicated by the nature of the post-translational modifications added to the C-terminal tails and thus determined by a combination of several factors at the C-termini requiring further analysis.

Rabs control essential endosomal functions including the regulation of phosphoinositide metabolism, vesicle transport and fusion, cargo sorting, and the recruitment of signaling molecules, as well as the identity and connectivity of the different classes of organelles ^17,70,71^. As such they are not only regulators of membrane trafficking, but indirectly also control cell signaling, polarity, migration and division. Consequently, the extensive Rab degradation induced by MCF could perturb many cellular processes. Nearly one-third of all Rabs associate with the Golgi and are important for the recruitment of effectors including tethering complexes, scaffolding proteins, kinases, and molecular motors required to maintain its architecture ^72^. Rab dysfunction interferes with the proper recruitment of these Golgi related proteins and results in disintegration ^73^. In fact, disruption of Rab1B homeostasis, or inactivation of just Rab1A or Rab1B in tissue culture cells affects Golgi stability and leads to its fragmentation ^19,29,35^. Approximately 38% (18/48) of the Rabs tested in the screen have been found to localize with the Golgi according to the UniProt database. Of those, 50% (9/18) are degraded or cleaved when co-expressed with MCF and would thus result in the extensive vesiculation of the Golgi caused by MCF ^6,8^. Furthermore, there is increasing evidence indicating Rabs localize to and play a role in maintaining mitochondrial fitness ^74-77^. As such it is possible Rab degradation also causes the mitochondrial fragmentation and induction of mitochondrial-dependent apoptosis previously observed to be caused by MCF in cells ^9^.

Fragmentation of the Golgi has been shown to increase the number of LC3 puncta and induce autophagy by decreasing the mTOR related signaling that inhibits autophagy and increasing the supply for autophagosomal biogenesis ^78,79^. Furthermore, Rabs play a role in several steps of autophagy ^80^. Loss of Golgi-associated proteins recruited by Rabs results in aberrant autophagy ^79^. C-terminal truncations of Rab4 increase the formation of autophagic structures and knockdowns of Rab5 inhibit mTOR activity ^81-83^. Knockouts of Rab1B result in the formation of significantly more LC3 puncta in tissue culture cells ^18^. However, the role of Rabs is not entirely clear as knockdowns and truncations have led to both upregulation and downregulation of autophagy activation ^18-21,29,84^. Interestingly, ectopic expression of MCF increases the number of autophagolysosomes in cells ^8^.

The significance of the induction of autophagy during infection is typically a host response to remove intracellular pathogens and to regulate inflammation and the immune response. How MCF modulates autophagy to benefit an extracellular pathogen is not yet clear. It is possible MCF dysregulates Rabs to inhibit maturation of phagolysosomes to suppress signaling and limit host detection of *V. vulnificus*. Knockouts of Rab1A or Rab1B are defective in protein secretion ^29,85,86^. Alternatively, MCF could stimulate autophagy to increase inflammation and tissue destruction at the site of infection to more easily cross protective barriers and disseminate.

Of significance, MCF is delivered to cells from an effector module that also delivers other protein effectors, including ABH. ABH cleaves phosphatidylinositol-3-phosphate to block recruitment of the complex required for membranes to form autophagosomes and thus inhibits initiation of autophagy ^87,88^. Therefore, the coordinated delivery of MCF with ABH as the ABH-MCF module could direct the loss of Golgi integrity and vesicle trafficking while inhibiting a host autophagy response. This would benefit the bacterium by arranging for cell death to occur in the absence of host signaling. Furthermore, we hypothesize the tethering allows ABH to direct MCF to membranes and thereby coordinates activation of MCF until it has reached its optimal location. Transfections with preprocessed aMCF, show deficient Rab degradation and cleavage, and suggest localization to membranes to increase proximity to Rabs is required for efficient activity (Fig. 1d and Supplementary Fig. 4a).

Interestingly, in addition to ABH and MCF, several *V. vulnificus* MARTX toxin effectors also either activate, target, or alter the downstream signaling of host GTPases. The Domain X (DmX) effector binds ARFs, the Rho GTPase-inactivation domain (RID) fatty acylates Ras homology (Rho) family members, and the Ras/Rap1-Specific Endopeptidase (RRSP) effector directly cleaves Rat sarcoma (Ras) and Ras-related protein 1 (Rap1) ^6,16,89-91^. This is an intriguing adaptation for an extracellular pathogen as typically intracellular pathogens alter host GTPases to increase survival inside host cells. Future studies examining how these MARTX toxin effectors coordinate to exploit host GTPase signaling will be vital to understanding how deadly *V. vulnificus* infections progress.

## Material and Methods

### Tissue culture

HEK 293T and Cos7 cells were cultured in Dulbecco’s modified Eagle’s medium (DMEM) (ThermoFisher 11965118) supplemented with 10% heat inactivated fetal bovine serum (FBS) (GeminiBio 900-108) and 1% penicillin-streptomycin at 37°C in 5% CO_2_. Cells were seeded in 100 mm tissue culture dishes or 12 -well tissue culture plates and grown to 60 - 70 % confluency.

### Bacterial strains and plasmids

A library of 48 fluorescently-tagged Rabs was obtained through previous collaborations as generated in Rzomp *et al.*, Heo and Meyer, Alvarez *et al.*, Smith *et al.*, Heo *et al.*, Sun *et al.*, Feng *et al.*, Barbero *et al.*, and Munafó and Colombo (Supplementary Table 1) ^24,92-99^. Rab isoforms were cloned into the peGFP-C2, peGFP-C3, peCFP-C1, peGFP-C1, or peGFP-N1 vector. N-terminal FLAG and C-terminal hemagglutinin (HA)-tagged vectors MCF^CA^, aMCF^CA^, MCF, aMCF, and MCF-eGFP plasmids were generated as described in Agarwal *et al*. ^7^. His-tagged vectors MCF^CA^, aMCF^CA^, MCF, aMCF, and ARF1 for protein purification were previously generated as described in Herrera *et al.* ^8^. GST-tagged Rab11A, 5A, 4A, 4B and peGFP-C3 Rab4B were generously provided by Seema Mattoo of Purdue University ^100^. Recombinant ACD (LF_N_ACD) was generated as previously described in Cordero *et al.* ^101^. CMCP6 *ΔvvhA rtxA1::bla* and CMCP6 *ΔvvhA rtxA1::mcf-bla V. vulnificus* strains were constructed as detailed in Agarwal *et al.* ^7^. gBlocks containing aMCF^domain^ ^II^(residues 85-324), aMCF^domain^ ^I^ (residues 1-84), Rab1B, Rab23, Rab2A, Rab6C, and Rab35 were synthesized by Integrated DNA Technologies (IDT) and cloned into pMCSG7, pGEX-KG, or peGFP-C2 expressions vectors using Gibson assembly (Supplementary Table 2).

### Rab phylogeny

Amino acid sequences for each Rab isoform were automatically aligned with ClustalW and manually inspected in MacVector v. 18.2.5 (43). An unrooted phylogenetic tree with branches transformed into a cladogram was visualized in FigTree v 1.4.4.

### Transfections and western blotting

HEK 293T cells were transfected using 1:3 μg of plasmid DNA to μL of 1 mg/mL polyethylenimine for ectopic gene expression; 50 μg of DNA per 100 mm tissue culture dish, and 3.1 μg per well in a 12 -well plate. After four hours the media was removed and fresh media was added, followed by incubation for an additional 14 hours.

For proteosome inhibition experiments cells were transfected as described above except the media was exchanged for fresh media after seven hours. Cells were then treated with 5 μM MG-132 (Millipore Sigma 474790) or dimethylsulfoxide (DMSO) vehicle control and incubated for 11 hours. For experiments inhibiting caspase activation, media was exchanged four hours post-transfection, cells were treated with 5 μM Z-VAD-FMK (Promega G7231) or DMSO vehicle control, and were incubated another 14 hours.

Following transfection, cells were collected, resuspended in lysis buffer (300 mM NaCl, 20 mM Tris–HCl pH 8, 1.1% Triton X-100) supplemented with ethylenediaminetetraacetic acid (EDTA) free protease inhibitor tablets (Thermo Scientific A32965), sonicated, and incubated for one hour at 4°C to recover whole cell lysates. 35 or 45 μg of total protein (same concentration for all samples in an individual gel) in the whole cell lysate was boiled for 10 min in 6× SDS loading buffer, separated by SDS-PAGE, and transferred to nitrocellulose membrane with the BioRad Mini TransBlot System. Western blots were then completed using anti-G/CFP (Invitrogen MA5-15256), anti-GST (Invitrogen A-5800), anti-HA (Sigma H6908), anti-α-Tubulin (Sigma T6074), anti-MCF (produced with purified MCF protein by Lampire Biological Laboratories, Pipersville PA), anti-ACD (produced with purified ACD protein by Lampire Biological Laboratories, Pipersville PA), anti-ARF1 (Novus Biologicals NBP1-97935), anti-mouse Immunoglobulin G (IgG) (Li-Cor 926-32210 or 926-68070), or anti-rabbit IgG (Li-Cor 926-32211 or 926-68071) antibodies and visualized using the Li-Cor Odyssey Fc imaging system.

### Rab degradation analysis

Each fluorescently-tagged Rab isoform in the screen was ectopically co-expressed with either MCF or the empty vector (p3xFlag-CMV-7.1) in three independent experiments. Proteins in lysates recovered for all three experiments were then separated in parallel on the same gel for comparison. The empty vectors the Rab isoforms are expressed in (peGFP-C2, pEGFP-C3, peCFP-C1, peGFP-C1, and pEGFP-N1) were also ectopically co-expressed with MCF or p3xFlag-CMV-7.1 in triplicate on the same gel. The amount of GFP-tagged Rab and tubulin in each lane for these gels was assessed by densitometry using NIH FIJI 3 imaging software v 2.3.0/1.53q, build d544a3f481. The degradation factor (DF) was calculated by the following formula where *d_MCF+_* represents density of the GFP or CFP band for samples co-expressed with MCF and *d_EV_* represents GFP or CFP band for samples co-expressed with the empty vector used to express MCF in the same set. *t_condition1_*is the density of the tubulin band of the Rab co-transfected with the empty vector in the first set all tubulin bands in the same blot will be normalized to.

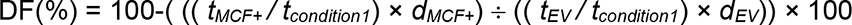

Where 100% DF is complete degradation. The DF was similarly determined for each empty fluorescent tag expression vector. Rab isoforms with an average DF 115% greater than the standard deviation of the mean of its respective empty vector were determined to be significantly degraded due to MCF co-expression. Cleavage of Rabs was manually assessed for each isoform in the screen.

### Bacterial intoxications

HEK 293T cells were incubated with either *V. vulnificus* strain CMCP6 *ΔvvhA rtxA1::bla* (no effectors) or CMCP6 *ΔvvhA rtxA1::mcf-bla* (MCF only). Bacteria were grown to mid-log phase in Luria broth, collected by centrifugation and resuspended in phosphate-buffered saline (PBS). HEK 293T cells at 60 - 70% confluency were washed with PBS and fresh DMEM without antibiotics or FBS added, after which cells were treated with the indicated strain at an MOI of 5 for either 1.5 or 2.5 hours at 37°C with 5% CO_2_. The media and bacteria were then aspirated and whole cell lysates were collected as above and 40 μL of lysate from each intoxication was separated by SDS-PAGE. A band containing proteins between 20 - 30 kDa in size, corresponding to the expected mobility of Rab isoforms, was excised for each intoxication and analyzed by mass spectrometry analysis at the Northwestern University Proteomics Center of Excellence Core Facility. The number of peptides of each Rab isoform identified was normalized to the number of peptides identified of the housekeeping gene HPRT in each sample.

### Protein production

MCF constructs and ARF1 were purified as previously described in Herrera *et al.* ^8^. aMCF^domain^ ^II^ was lysed in Buffer A (500 mM NaCl, 10 mM Tris pH 8.0) with 3M guanidine hydrochloride and purified as the other MCF constructs in this buffer. Following collection the protein was dialyzed in Buffer A with 1M guanidine hydrochloride, then dialyzed with Buffer A containing 5 - 20% glycerol, and finally buffer exchanged into Buffer A. GST-tagged Rab isoforms were expressed, grown, and lysate collected as above and batch purified using glutathione sepharose (GE 17-0756-01). ACD (LF_N_ACD) was purified as described in Cordero *et al.* ^101^.

### Immunoprecipitation

To recover tagged protein from cells, whole cell lysate (acquired as described above) from transfections was incubated overnight at 4°C with mouse IgG agarose (Sigma A0919). The agarose was then removed by centrifugation and the lysate recovered and incubated overnight at 4°C with either anti-G/CFP monoclonal antibody agarose (MBL International D153-8) or EZview™ Red Anti-HA Affinity Gel (Sigma E6779). The beads were subsequently washed four times with lysis buffer supplemented with EDTA-free protease inhibitor tablets, washed once with 10 mM Tris–HCl pH 8.0, and protein eluted with 2 M NaSCN in 50 mM Tris–HCl pH 8.0, 150 mM NaCl.

CFP-Rab23 recovered from cell lysates by anti-G/CFP bead immunoprecipitation was separated by SDS-PAGE and gel bands corresponding to full-length and cleaved CFP-Rab23 were individually excised and analyzed by mass spectrometry analysis at the Northwestern University Proteomics Center of Excellence Core Facility using a no-enzyme search (Supplementary Fig. 8). CFP-Rab23 similarly recovered from cell lysates was also analyzed by two-dimensional western analysis as previously described ^8^.

For co-immunoprecipitation assays with purified recombinant proteins, variations of 2.5 μM MCF constructs, 4 μM GST-Rab isoforms, 4 μM ARF1, and 2.5 μM ACD were incubated in RAX buffer (20 mM Tris pH 7.5, 150 mM NaCl, 10 mM MgCl_2_) overnight (or for two hours prior to overnight as indicated) at 37°C. Anti-GST pull-downs were completed with these protein suspensions subsequently incubated with Pierce™ Glutathione Agarose (ThermoFisher 16100) for an hour at 4°C in binding buffer (PBS pH 7.0 with 1 mM Tris (2-carboxyethyl) phosphine), beads then washed six times with binding buffer, and bound protein eluted with 50 mM Tris-HCl pH 8.0 with 10 mM glutathione.

### Fluorescence Microscopy

Immunofluorescence microscopy images were taken at the Northwestern University Center for Advanced Microscopy using the Nikon A1R+ GaAsP Confocal Laser Microscope. Following transfection Cos7 cells grown on glass coverslips were washed, fixed, permeabilized, and blocked as previously described in Herrera *et al.* prior to staining with anti-Rab1B (Thermo Scientific 17824-1-AP) in staining buffer (1% bovine serum albumin (BSA) plus 0.05% Tween 20 in PBS) overnight ^8^. Coverslips were washed and incubated with secondary goat anti-rabbit Alexa Fluor 647 antibody (ThermoFisher A-21244) for detection of Rab1B. Images were taken in at least three fields of cells ectopically expressing MCF or the empty vector, as identified by GFP fluorescence.

### Crystallization, data collection, and structure determination

aMCF^CA^ in *E. coli* (DE3) BL21 (pMAGIC) ^102,103^ was cultured in enriched M9 SeMET High-Yield Growth Media (Medicilon MD045004) supplemented with kanamycin and ampicillin and grown to an OD_600_ of 1.6 - 1.8 at 37°C ^104^. 0.16 g/L of selenomethionine and 0.5 mM isopropyl β-D-1-thiogalactopyranoside (IPTG) was then added to the cells and they were grown for an additional 16 hours at 25°C. Cells were harvested by centrifugation, resuspended in Buffer A+ (10 mM Tris-HCl pH 8.3, 500 mM NaCl, 10% glycerol, 0.1 % IGEPAL® CA-630 (MilliporeSigma I3021) and protease inhibitor tablets), and frozen at -20°C. Cells were then thawed, sonicated for 20 minutes at 40% amplitude (pulse 5 seconds on and 10 seconds off), and centrifuged. aMCF^CA^ was then purified from the supernatant using the AKTA protein purification system through a Ni-NTA HisTrap affinity column (GE Healthcare GE17-5248-02) with loading buffer (10 mM Tris-HCl pH 8.5, 500 mM NaCl, 1 mM Tris (2-carboxyethyl) phosphine (TCEP)), washing buffer (10 mM Tris-HCl pH 8.5, 500 mM NaCl, and 25 mM imidazole), and elution buffer (10 mM Tris pH 8.3, 500 mM NaCl, and 500 mM imidazole). The eluted protein was loaded onto a HiLoad Superdex 26/600 column for size exclusion chromatography. The His-tag was cleaved from the protein by incubation with the tobacco etch virus (TEV) protease, and an additional Ni-NTA HisTrap affinity chromatography purification was performed to remove the protease, uncut protein, and the affinity tag from aMCF^CA^ ^105^. Fractions were concentrated separately at 8.5 mg/mL.

Crystallization was set up using the sitting drop method with a 1:1 protein to reservoir ratio and a drop volume of 2 μL with the commercial ammonium sulfate suite screen (NeXtal 130905). Crystals grew from condition 15 (0.2 M cesium sulfate in 2.2 M ammonium sulfate) and were cryoprotected in 25% sucrose in 3.6 M ammonium sulfate. Diffraction quality crystals were screened and data collected at the 21ID-G beamline of the Life Sciences–Collaborative Access Team (LS-CAT) at the Advanced Photon Source (APS), Argonne National Laboratory. Images were indexed, integrated, and scaled using HKL-3000 (Supplementary Table 3) ^105^. The structure of aMCF^CA^ was determined by molecular replacement using the structure of aMCF^CS^ in complex with ARF3^Q71L^ (PDB code 6ii6) as a search model. The model was split into two domains and automated multiple component search was used in PHASER ^105^ from the CCP4 suite ^105^. The initial solution went through several cycles of restrained refinement in REFMAC v. 5.8.0257 ^105^, and a linker region was manually built in COOT ^105^. Several additional rounds of refinement were completed with REFMAC and manual corrections performed using COOT prior to generating water molecules in ARP/wARP ^105^ and fitting ligands into maps in COOT. Translation/Libration/Screw (TLS) groups were generated using the TLSMD server (http://skuldbmsc.washington.edu/~tlsmd/) and TLS corrections applied at the final steps of model refinement ^105^. The final structure was validated using MolProbity (http://molprobity.biochem.duke.edu/). Coordinates of the model and experimental data were deposited to the Protein Data Bank (https://www.rcsb.org/) with the assigned PDB code 8SFG^105^.

### Structure prediction and model generation

Structure prediction models were generated with the structure prediction tool in ChimeraX v 1.5.dev202207070159 using ColabFold v1.5.1 on the Google Colab virtual machine or with Alphafold2 v 2.2.2 (max template date 2022-05-23) on Northwestern’s Structural Biology Facility cluster. All structures were visualized and analyzed in ChimeraX. Structural alignment of models was completed using the matchmaker tool in ChimeraX. The published solved structure of aMCF^CS^-ARF3^Q71L^ (PDB code 6ii6), aMCF^CS^ (PDB code 6ii0) and Rab1B (PDB code 4i1o) were used for comparisons ^6,26^. The end of the fifth alpha helix in Rab isoforms was manually determined from the prediction models, allowing for breaks in the helix no larger than 6 amino acids. Any residues past the determined fifth alpha helix were designated as the hypervariable domain “tail” for the respective Rab. The complex structure of each Rab isoform in our screen with aMCF was individually predicted. These complexes were aligned to aMCF^CS^-ARF3^Q71L^ (PDB code 6ii6) using matchmaker and residues in Rabs whose spatial orientation aligned with the residues in ARF3^Q71L^ previously characterized to be important for binding were identified ^6^.

## Acknowledgements

This work was supported by funding from the NIAID grants R01 AI092825-09 (to K.J.F.S.) and K99 GM143571-01 (to A.H.). This project has been funded in part with Federal funds from the Department of Health and Human Services, National Institutes of Health, and National Institute of Allergy and Infectious Diseases under contract no. HHSN272201700060C and 75N93022C00035 (to K.J.F.S). This research used resources of the Advanced Photon Source, a U.S. Department of Energy (DOE) Office of Science User Facility operated for the DOE Office of Science by Argonne National Laboratory under contract no. DE-AC02- 06CH11357. Use of the LS-CAT Sector 21 was supported by the Michigan Economic Development Corporation and the Michigan Technology Tri-Corridor (grant 085P1000817). Access to LS-CAT is a service of the Northwestern University Structure Biology Facility, which is generously supported by NCI CCSG P30 CA060553. This project includes work performed at the Northwestern University Center for Advanced Microscopy, Structural Biology Facility, and the Proteomics Core, all generously supported by NCI CCSG P30 CA060553 awarded to the Robert H Lurie Comprehensive Cancer Center. Proteomics services are further supported by an instrumentation award (S10OD025194) from NIH Office of Director, and the National Resource for Translational and Developmental Proteomics supported by P41 GM108569. Molecular graphics and analyses performed with UCSF ChimeraX, developed by the Resource for Biocomputing, Visualization, and Informatics at the University of California, San Francisco, with support from National Institutes of Health R01-GM129325 and the Office of Cyber Infrastructure and Computational Biology, National Institute of Allergy and Infectious Diseases.

We thank Dr. Seema Mattoo for sharing Rab expression plasmids, Dr. Suzanne Pfeffer for feedback on development of the project, and Jason A. Pattie for his help running Alphafold2 on Northwestern’s Structural Biology Facility cluster.

## Author contributions

A.H. designed, conducted, and analyzed all experiments, drafted and edited the manuscript, and provided funding. M.P. conducted experiments. M.R.L. and G. M. conducted protein purification and structure determination and contributed to data analysis and editing of the manuscript, J.B contributed reagents and assisted in experimental design, K.J.F.S. contributed to experimental design and analysis, edited the manuscript, and provided funding.

## Data availability

Coordinates for the x-ray structure of MCF has been deposited to the RCSB Protein Data Bank with the code 8SFG. All other data are found within the manuscript or supplemental data. Expression plasmids are available upon request from.

