## Supplemental Data for "*Vibrio* MARTX toxin processing and degradation of cellular Rab GTPases by the cytotoxic effector Makes Caterpillars Floppy"

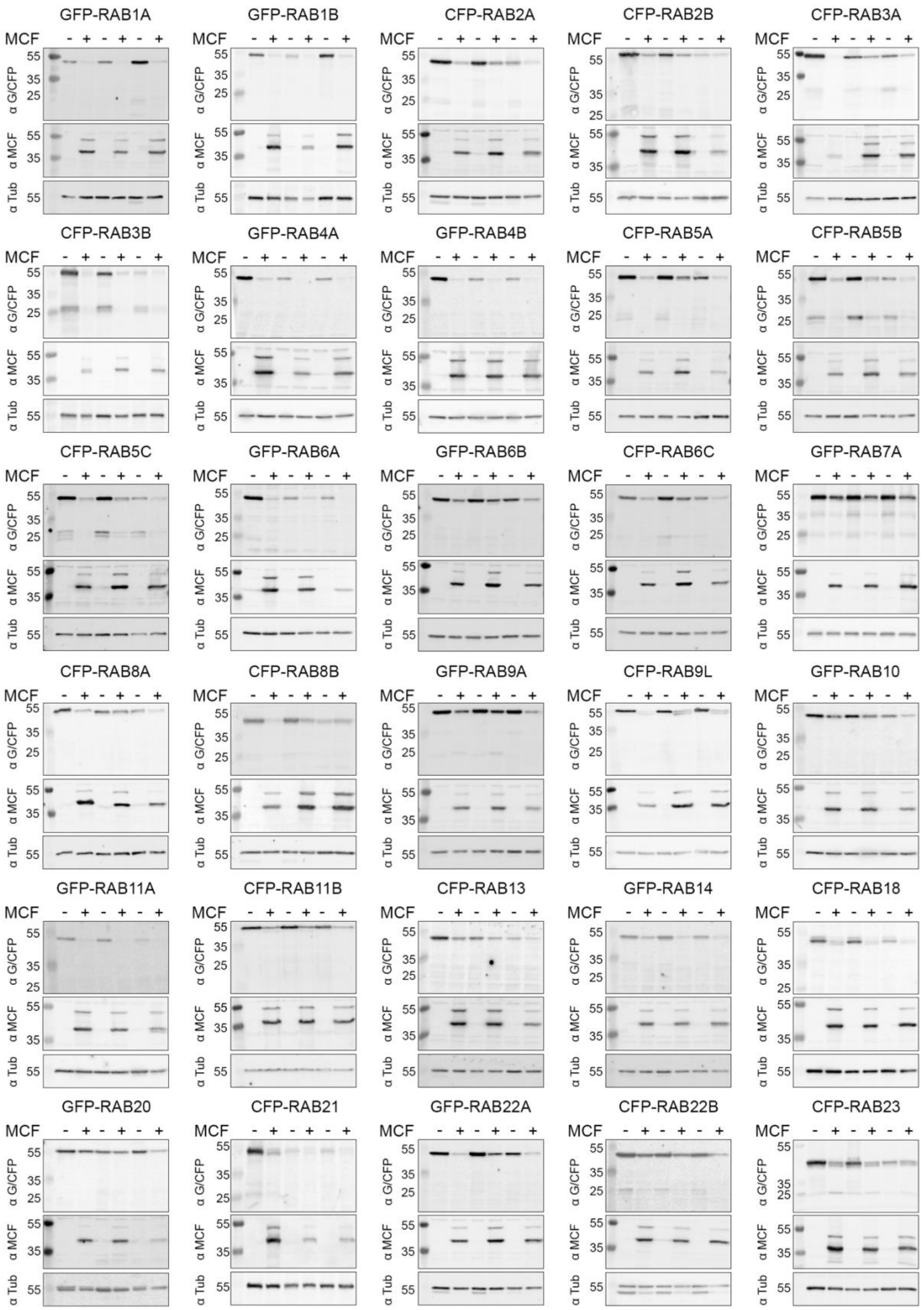
GFP-RAB1A
GFP-RAB1B
CFP-RAB2A
CFP-RAB2B
CFP-RAB3A
CFP-RAB3B
GFP-RAB4A
GFP-RAB4B
CFP-RAB5A
CFP-RAB5B
CFP-RAB5C
GFP-RAB6A
GFP-RAB6B
CFP-RAB6C
GFP-RAB7A
CFP-RAB8A
CFP-RAB8B
GFP-RAB9A
CFP-RAB9L
GFP-RAB10
GFP-RAB11A
CFP-RAB11B
CFP-RAB13
GFP-RAB14
CFP-RAB18
GFP-RAB20
CFP-RAB21
GFP-RAB22A
CFP-RAB22B
CFP-RAB23
MCF - + - + - +
α G/CFP
α MCF
α Tub
55
35
25

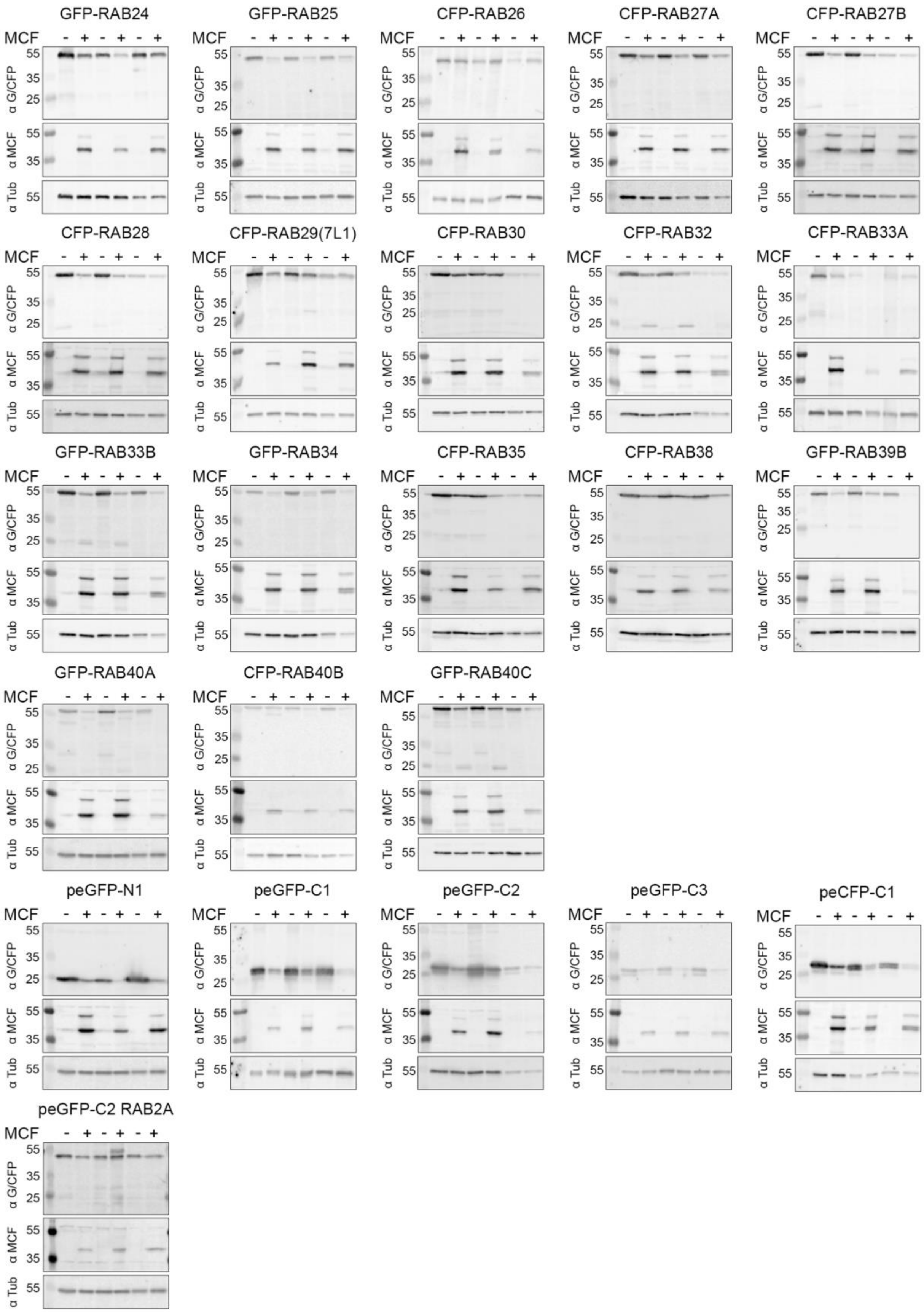
GFP-RAB24
GFP-RAB25
CFP-RAB26
CFP-RAB27A
CFP-RAB27B
CFP-RAB28
CFP-RAB29(7L1)
CFP-RAB30
CFP-RAB32
CFP-RAB33A
GFP-RAB33B
GFP-RAB34
CFP-RAB35
CFP-RAB38
GFP-RAB39B
GFP-RAB40A
CFP-RAB40B
GFP-RAB40C
peGFP-N1
peGFP-C1
peGFP-C2
peGFP-C3
peCFP-C1
peGFP-C2 RAB2A
MCF - + - + - +
α G/CFP
α MCF
α Tub
55
35
25

Supplementary Figure 1. The MCF toxin causes variable degradation of 48 fluorescently-tagged Rabs when co-expressed. Fluorescently tagged Rabs were co-transfected with MCF (or an empty vector control, p3xFlag-CMV-7.1) in HEK 293T cells in three independent experiments. The vector each Rab is expressed from is indicated in Supplementary Table 1, unless directly labelled on blot. Empty fluorescently tagged vectors used for Rab expression were also co-transfected with MCF or p3xFlag-CMV-7.1 in three independent experiments. Total protein (45 or 35 μg; same within each blot) from the triplicate experiments was analyzed by western blot using whole cell lysates with anti- GFP/CFP (α G/CFP), anti MCF (α MCF), and anti-α-Tubulin (α Tub) antibodies.

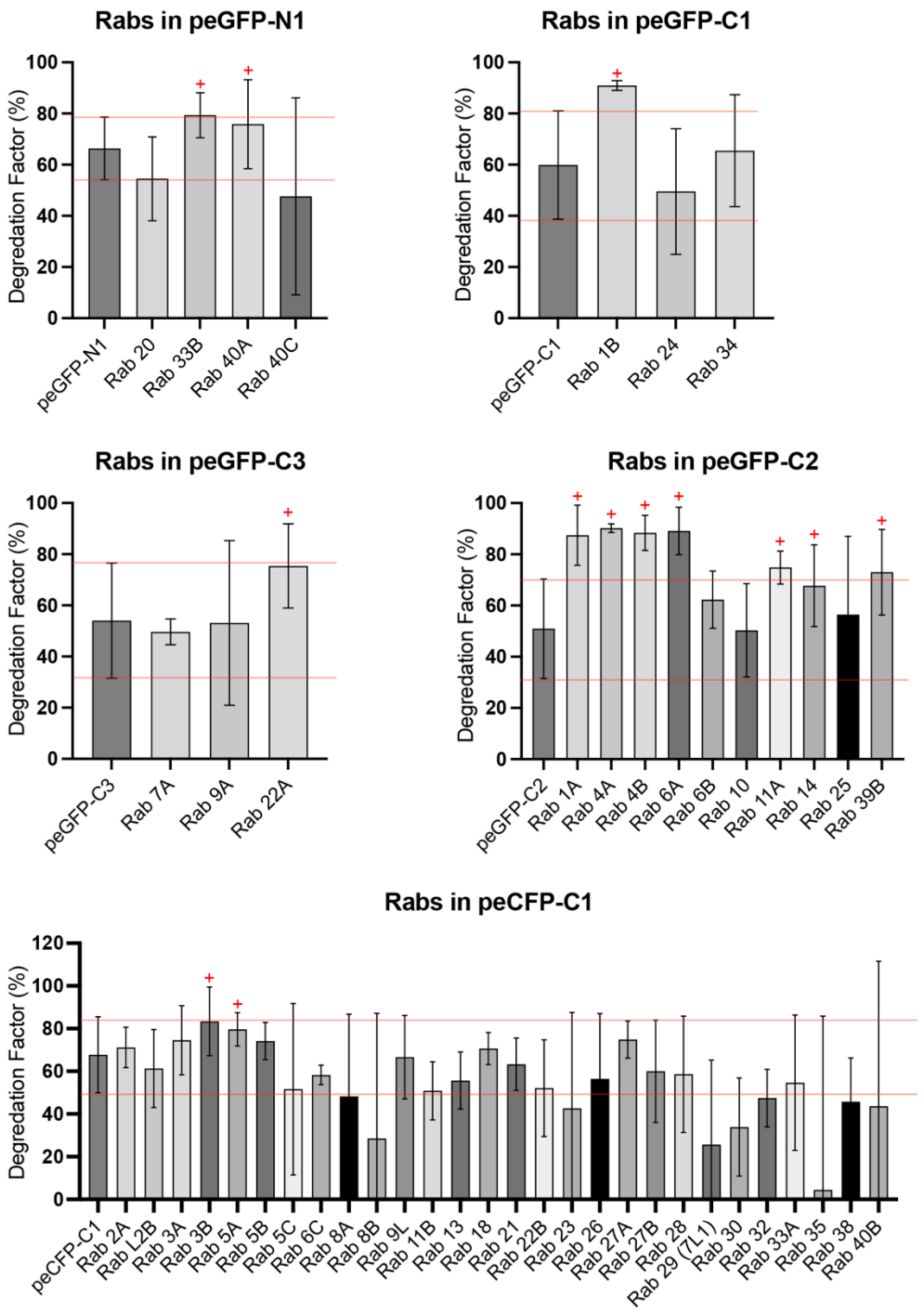


Supplementary Figure 2. Degradation factor (DF) of Rabs co-expressed with MCF compared to that of the empty vector each is expressed from . Densitometry of bands on western blots (Supplementary Fig. 1) from three independent experiments was used to determine the DF of each Rab, where 100% is complete degradation (see methods). The DF of each replicate was plotted onto a bar graph showing the mean with error bars for standard deviation (SD). Each Rab isoform is plotted in the same graph as the empty vector (peGFP C2, pEGFP-C3, peCFP-C1, peGFP-C1, or pEGFP-N1) it is expressed from. The SD limits of the DF for each empty vector is drawn across the blot in red. Rabs with a DF 115% greater than the SD of their respective empty vector are denoted with a "+".

a

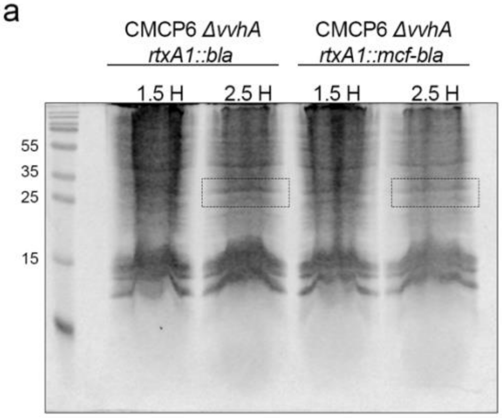


b

| # of Peptides Identified 2.5 H Post Intoxication | | | |
|---|---|---|---|
| Rab | CMCP6 *rtxA1::bla* | CMCP6 *rtxA1::mcf-bla* | Screen Results |
| Rab7A | 4 | 1 | unaffected |
| Rab1A | 2 | 3 | degraded |
| Rab5B | 4 | 1 | cleaved |
| Rab14 | 3 | 0 | degraded |
| Rab21 | 9 | 0 | cleaved |
| Rab11B | 2 | 0 | cleaved |
| HPRT | 14 | 10 | N/A |
| ATP5PB | 15 | 8 | N/A |

c

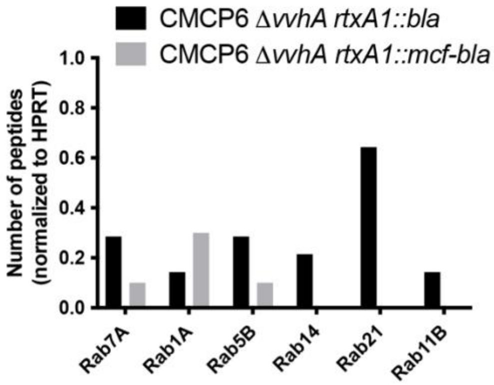


Supplementary Figure 3. Natural MARTX toxin delivery of the MCF effector during bacterial intoxication reduces the number of Rab peptides recovered from tissue culture cells. **a** HEK 293T cells were intoxicated with *V. vulnificus* with either an effectorless (CMCP6 *ΔvvhA rtxA1::bla*) or MCF only (CMCP6 *ΔvvhA rtxA1::mcf-bla*) encoding MARTX toxin for 1.5 or 2.5 hours at an MOI of 5. 40 μL of whole cell lysate was analyzed by SDS-PAGE, and bands encompassing all proteins ranging from 30 - 20 kDa in size were excised (dotted boxes) for intoxications at 2.5 hours. **b** Table shows the total number of peptides recovered of each Rab, and housekeeping proteins hypoxanthine phosphoribosyltransferase (HPRT) and ATP synthase F (0) complex subunit B1 (ATP5PB) by mass spectrometry analysis of the excised bands in (**a**). **c** Graph of total number of peptides recovered in (**b**) of each Rab normalized to HPRT.

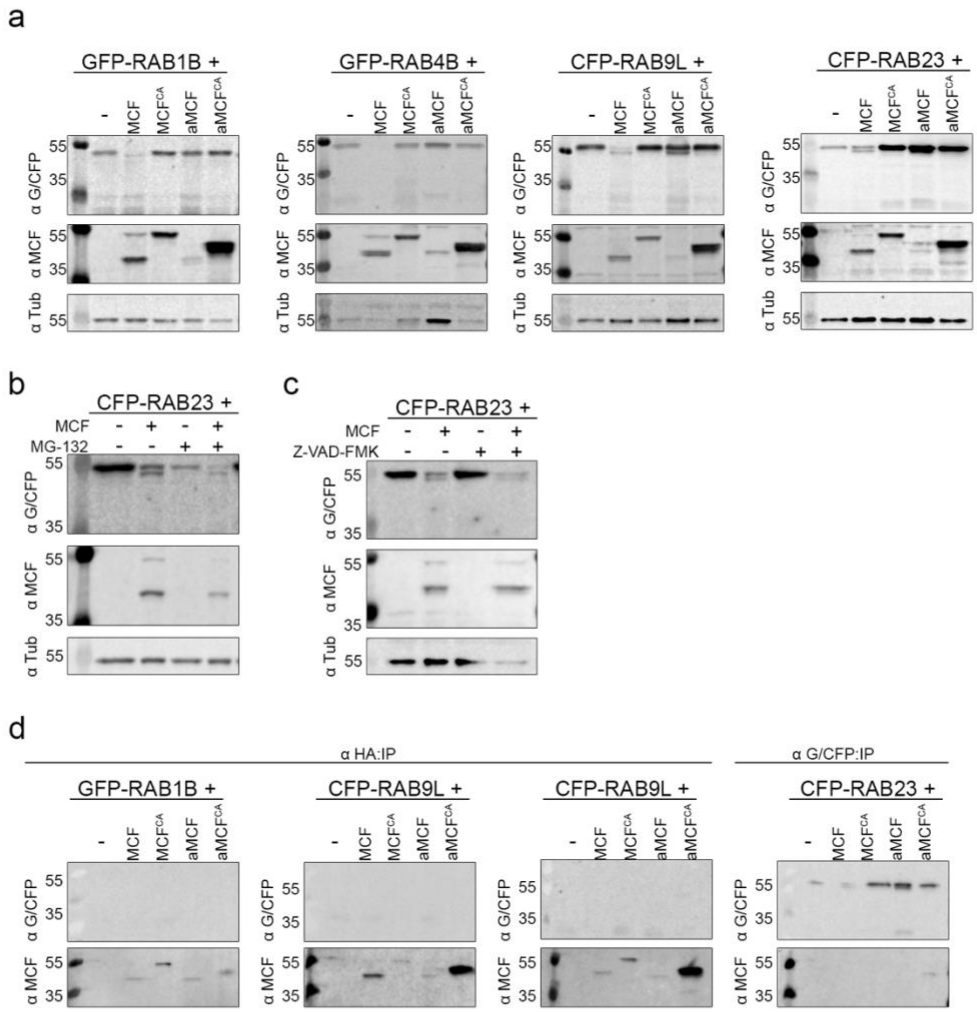


Supplementary Figure 4. Proteolytic activity of MCF is required for Rab degradation. **a** GFP-Rab1B, GFP-Rab4B, CFP-Rab9L, and CFP-Rab23 were co-expressed with MCF, catalytically inactive MCF (MCF$^{CA}$), autoprocessed MCF (aMCF), or catalytically inactive aMCF (aMCF$^{CA}$) in HEK 293T cells. Cell lysates recovered from these transfections were used to complete western blots with anti-GFP/CFP ($\alpha$ G/CFP), anti-MCF ($\alpha$ MCF), and anti-$\alpha$-Tubulin ($\alpha$ Tub) antibodies. **b, c** HEK 293T cells were co-transfected with MCF and CFP-Rab23 in the presence of 5 $\mu$M of either (**b**) MG-132 or (**c**) Z-VAD-FMK and western blots completed as in (**a**). **d** Co-expression experiments were completed with the MCF constructs used in (**a**) which have a hemagglutinin (HA)-tag at their C-terminus and GFP-Rab1B, CFP-Rab9L, or CFP-Rab23. Whole cell lysates recovered from the experiments were used to complete anti-HA or anti-G/CFP co-immunoprecipitation assays and western blots performed as above on eluted protein.

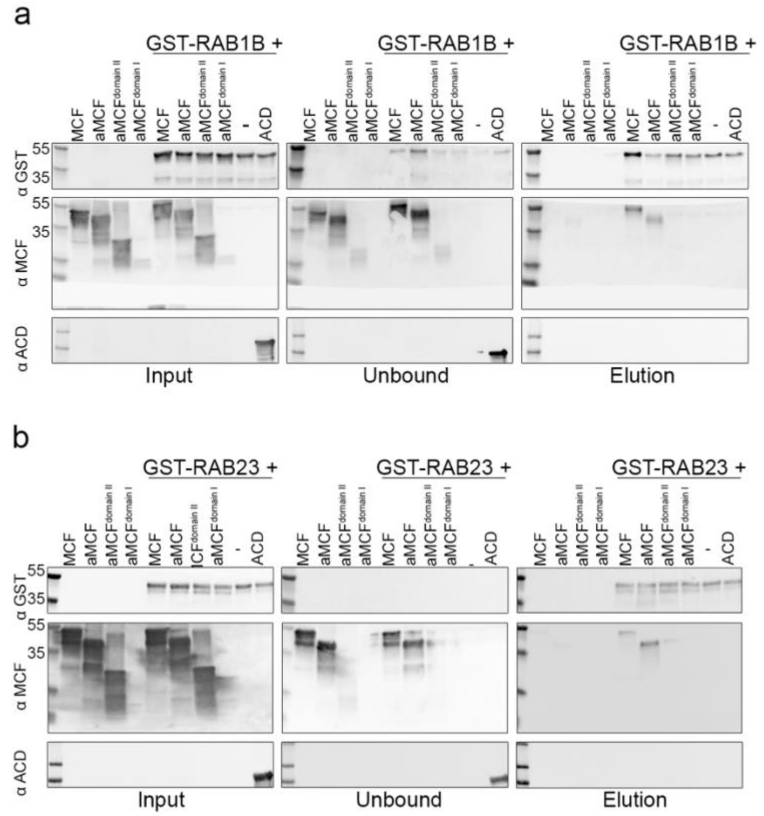


Supplementary Figure 5. Recombinant MCF directly binds to Rab1B and Rab23. **a, b** MCF, aMCF, residues 85-324 of aMCF ($aMCF^{domain\ II}$), or residues 1-84 of aMCF ($aMCF^{domain\ I}$) were incubated with purified (**a**) GST-Rab1B or (**b**) GST-Rab23 at 37°C overnight. The ACD was used as a control for non-specific binding. The incubated samples (input), flow through not bound to GST beads (unbound), and protein eluted from the beads following anti-GST immunoprecipitation (IP) (elution) were analyzed by western blot with anti-GST (α GST), anti-MCF (α MCF), and anti-ACD (α ACD) antibodies.

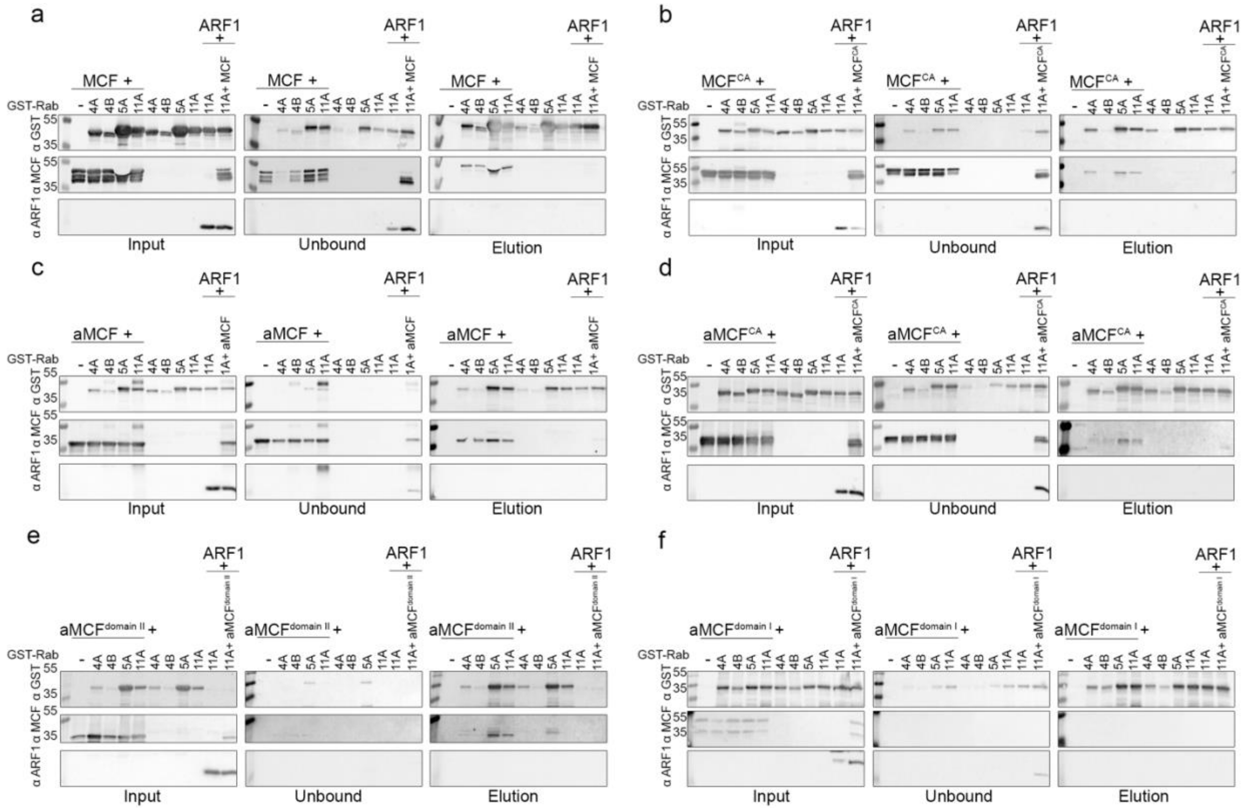


Supplementary Figure 6. Several recombinant Rab isoforms directly bind to MCF. **a-f** Purified GST-Rab4A, GST-Rab4B, GST-Rab5A, and GST-Rab11A were incubated with (**a**) MCF, (**b**) catalytically inactive MCF ($MCF^{CA}$), (**c**) autoprocessed MCF (aMCF), (**d**) catalytically inactive aMCF ($aMCF^{CA}$), (**e**) residues 85-324 of aMCF ($aMCF^{domain\ II}$), or (**f**) residues 1-84 of aMCF ($aMCF^{domain\ I}$) at 37°C overnight. One reaction for each MCF includes ARF1 added simultaneously with GST-Rab11A as indicated. The incubated samples (input), flow through not bound to GST beads (unbound), and protein eluted from the beads following anti-GST immunoprecipitation (IP) (elution) were analyzed by western blot with anti-GST (α GST), anti-MCF (α MCF), and anti-ARF1 antibodies (α ARF1) antibodies.

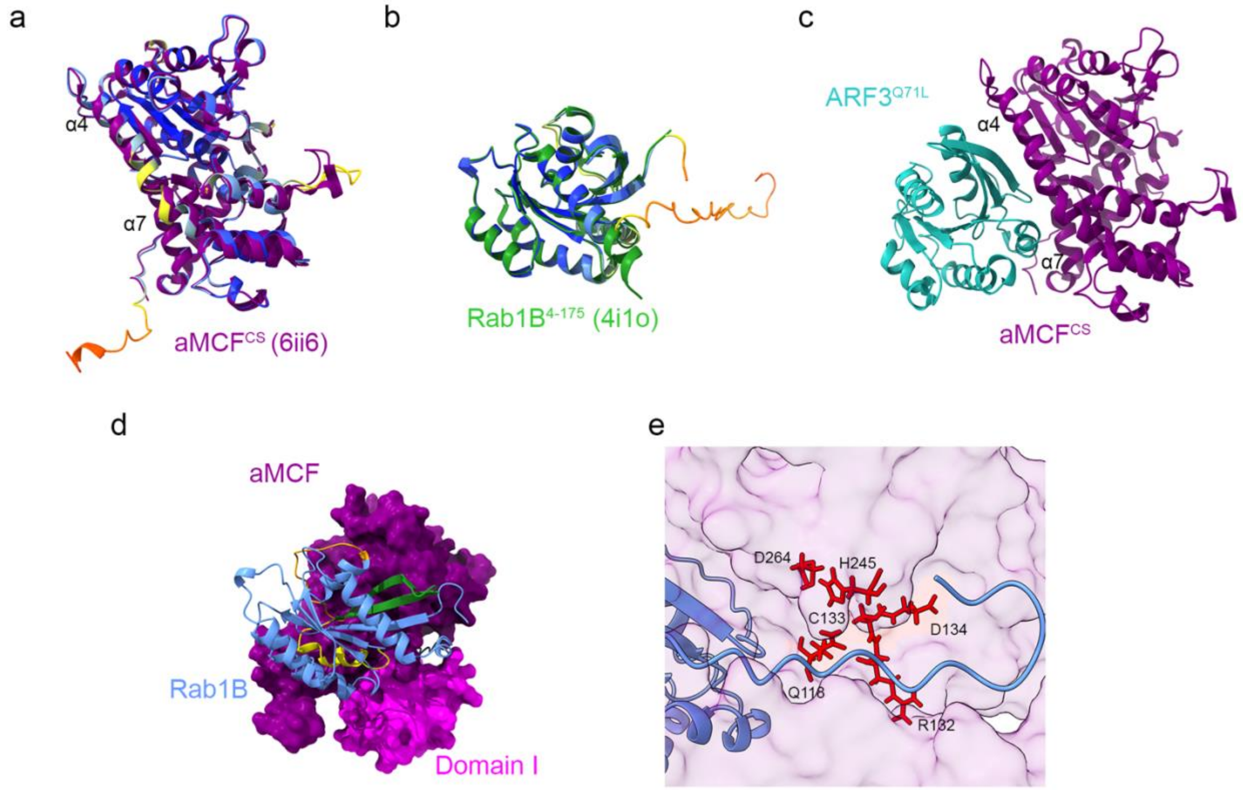


Supplementary Figure 7. Rabs are predicted to interact with autoprocessed MCF mostly through residues within their G domain. **a** The predicted ribbon structure of aMCF in the color bfactor palette Alphafold rainbow coloring according to pLDDT confidence overlayed on $aMCF^{CS}$ (extracted from the $ARF3^{Q71L}$-$MCS^{CS}$ complex; PDB code 6ii6) [1] (purple). **b** The predicted ribbon structure of Rab1B (extracted from the predicted Rab1B-aMCF structure) in the color bfactor palette Alphafold rainbow coloring according to pLDDT confidence overlayed on $Rab1B^{4\text{-}175}$ (extracted from the structure of *L. pneumophila* GAP domain of LepB in complex with Rab1B bound to GDP and BeF3 ; PDB code 4i1o) [2] (green). **c** Previously solved ribbon co-structure of ADP Ribosylation Factor 3 mutated to mimic the active state ($ARF3^{Q71L}$) (light sea green) with catalytically inactive aMCF ($aMCF^{CS}$) (purple) (PDB code 6ii6) [1]. **d** Predicted complex surface structure of aMCF, with its domain I, residues 1-84, ($aMCF^{domain\ I}$) highlighted in magenta, bound to the ribbon structure of Rab1B (cornflower blue), with its switch one (orange), interswitch (green), and switch two (yellow) regions specified. **e** Close up of channel formed from predicted complex of aMCF bound to Rab1B (from Fig. 3a), with residues important for its catalytic activity labelled.

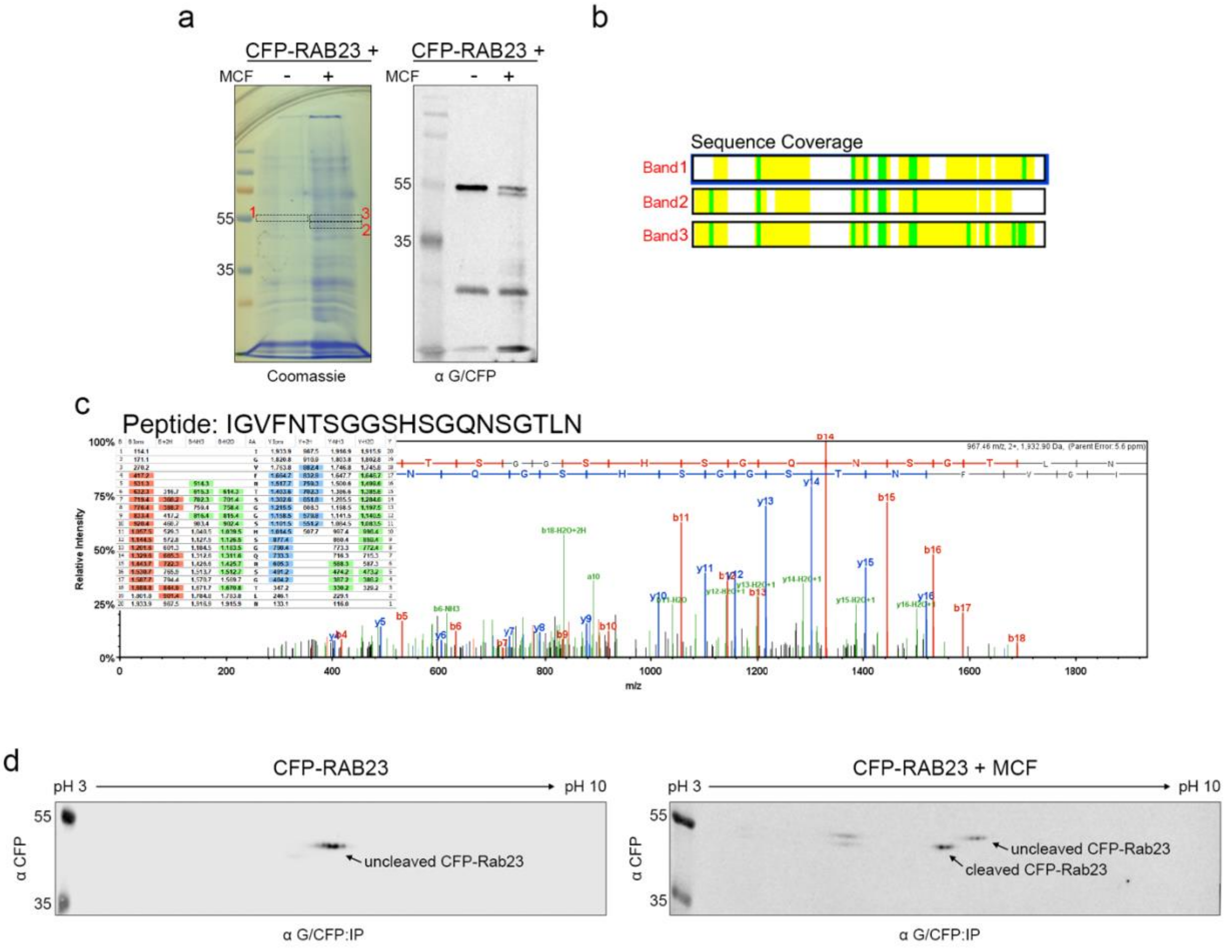


Supplementary Figure 8. MCF toxin causes only particular populations of post-translationally modified Rab 23 to be cleaved at their C-terminus when co-expressed. **a** Coomassie gel of CFP-Rab23 recovered from whole cell lysates of HEK 293T cells co-transfected with MCF or empty vector control p3xFlag-CMV-7.1 through anti-CFP immunoprecipitation (IP), with corresponding western blot, using anti-CFP antibodies, of same IP samples. Dashed-lined boxes indicate bands corresponding to uncleaved CFP-Rab23 (bands 1 and 3), and cleaved CFP-Rab23 (band 2). **b** Bands outlined in (**a**) were excised and analyzed by mass spectrometry. Schematic shows the sequence coverage for CFP-Rab23 in each band. **c** Spectrum and fragmentation table of the "IGVFNTSGGSHSGQNSGTLN" peptide in band 2. **d** Two-dimensional western analysis of CFP-Rab23 recovered from anti-CFP IPs of co-transfection experiments with MCF or p3xFlag-CMV-7.1 as in (**a**) on immobilized pH gradient

(IPG) strips pH 3-10. Gel shows populations of CFP-Rab23 present without (left) and with (right) MCF co-expression.

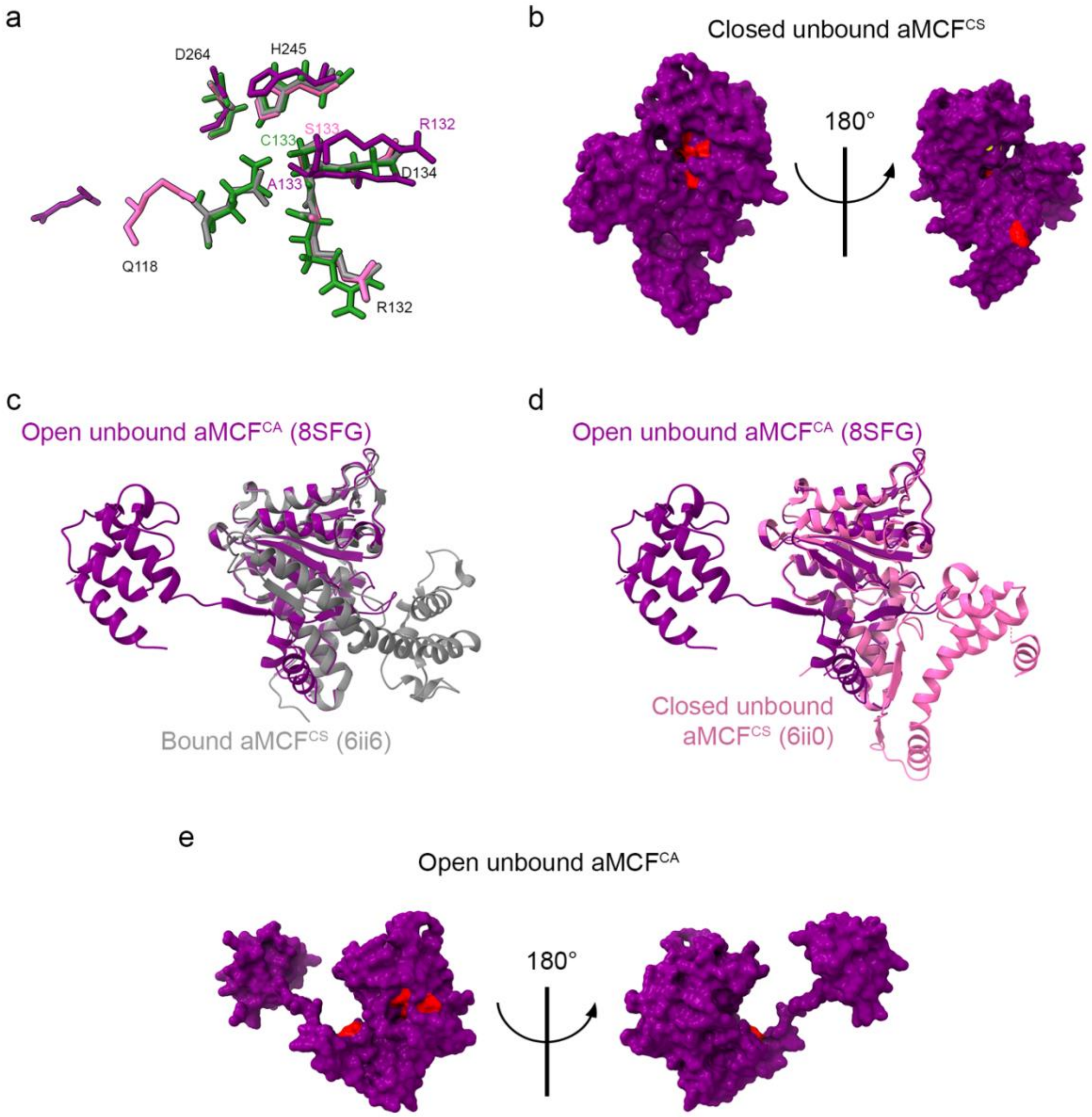


Supplementary Figure 9. The unbound autoprocessed MCF toxin transitions between two conformational states. **a** Overlay of residues for closed unbound aMCF$^{CS}$ (PDB code 6ii0) [1] (pink), open unbound aMCF$^{CA}$ (PDB code 8SFG) (purple), closed bound aMCF$^{CS}$ (derived from PDB code 6ii6) [1] (gray), and predicted aMCF (derived from complex prediction with Rab1B as in Fig. 3a) (green) important for its catalytic activity. **b** Two views of the surface structure of closed unbound aMCF$^{CS}$ (PDB code 6ii0) [1], with residues Arg132, Cys133, Asp134, His245, and Gln118 in red, and Asp264 in yellow. **c** Overlay of the ribbon structures of closed bound aMCF$^{CS}$ (gray) (PDB code 6ii6) [1] with open aMCF$^{CA}$ (purple) (PDB code 8SFG). **d** Overlay of the ribbon structures of closed unbound aMCF$^{CS}$ (pink) (PDB code 6ii0) [1] with open aMCF$^{CA}$ (purple) (PDB code 8SFG). **e** Two views of the surface structure of open unbound aMCF$^{CA}$ (PDB code 8SFG), with residues Arg132, Cys133, Asp134, His245, and Gln118 in red, and Asp264 in yellow.

.

| | | | | | | | | | | | | | | | | | | | | | | |
|---|---|---|---|---|---|---|---|---|---|---|---|---|---|---|---|---|---|---|---|---|---|---|
| ***ARF3*** | E17 | R19 | K38 | L39 | I46 | I49 | G50 | F51 | V53 | E54 | T55 | T64 | W66 | D72 | K73 | I74 | P76 | L77 | H80 | Y81 | Q83 | E115 |
| Rab4A | L13 | K15 | I34 | G35 | N43 | I46 | G47 | V48 | F50 | G51 | S52 | Q65 | W67 | E73 | R74 | F75 | S77 | V78 | S81 | Y82 | R84 | A115 |
| Rab4B | L8 | K10 | I29 | E30 | N38 | I41 | G42 | V43 | F45 | G46 | S47 | Q60 | W62 | E68 | R69 | F70 | S72 | T74 | S76 | Y77 | R79 | A110 |
| Rab14 | I11 | K13 | T32 | E33 | P41 | I44 | G45 | V46 | F48 | G49 | T50 | Q63 | W65 | E71 | R72 | F73 | A75 | V76 | S79 | Y80 | R82 | T113 |
| Rab6A | R12 | K15 | M34 | Y35 | Q43 | I46 | G47 | D49 | F50 | L51 | S52 | Q65 | W67 | E73 | R74 | F75 | S77 | L78 | S81 | Y82 | R84 | S117 |
| Rab22A | R4 | K7 | V26 | E27 | N35 | I38 | G39 | A40 | F42 | M43 | T44 | L57 | W59 | E65 | R66 | F67 | A69 | L70 | M73 | Y74 | R76 | G107 |
| Rab39B | Q8 | R10 | T29 | G30 | D38 | V41 | G42 | V43 | F45 | F46 | S47 | Q61 | W63 | E69 | R70 | F71 | S73 | I74 | A77 | Y78 | R80 | V111 |
| Rab11A | L11 | K13 | T32 | R33 | K41 | I44 | G45 | V46 | F48 | A49 | T50 | Q63 | W65 | E71 | R72 | Y73 | A75 | I76 | A79 | Y80 | R82 | A113 |
| Rab1B | L8 | K10 | A29 | D30 | I38 | I41 | G42 | V43 | F45 | K46 | I47 | Q60 | W62 | E68 | R69 | F70 | T72 | I73 | S76 | Y77 | R79 | A110 |
| Rab3B | M22 | K24 | A43 | D44 | V52 | V55 | G56 | I57 | F59 | K60 | V61 | Q74 | W76 | E82 | R83 | Y84 | T86 | I87 | A90 | Y91 | R93 | S124 |
| Rab33B | I33 | K35 | C54 | A55 | E63 | I66 | G67 | V68 | F70 | R71 | E72 | Q85 | W87 | E93 | R94 | F95 | K97 | S98 | H102 | Y103 | R105 | L136 |
| Rab1A | Y10 | L11 | A32 | D33 | I41 | I44 | G45 | V46 | F48 | K49 | I50 | Q63 | W65 | E68 | R72 | F73 | T75 | I76 | S79 | Y80 | R82 | A113 |
| Rab5A | Q20 | K22 | V41 | D30 | E50 | I53 | G54 | A55 | F57 | L58 | T59 | E72 | W74 | E80 | R81 | Y82 | S84 | L85 | M88 | Y89 | R91 | A122 |
| *Rab40A* | | | | | | | | | | | | | | | | | | | | | | |
| Rab25 | V12 | K14 | T33 | R34 | R42 | I45 | G46 | V47 | F49 | S50 | T51 | Q64 | W66 | E72 | R73 | Y74 | A76 | I77 | A80 | Y81 | R83 | A114 |
| Rab11B | L11 | K13 | T32 | R33 | K41 | I44 | G45 | V46 | F48 | A49 | T50 | Q63 | W65 | E71 | R72 | Y73 | A75 | I76 | A79 | Y80 | R82 | A113 |
| Rab23 | A9 | K11 | C30 | K31 | K39 | I42 | G43 | V44 | F46 | L47 | E48 | L60 | W63 | E69 | E70 | F71 | A73 | I74 | A77 | Y78 | R80 | G112 |
| Rab34 | I52 | L54 | C73 | K74 | K82 | I85 | G86 | V87 | F89 | E90 | M91 | Q104 | W106 | E112 | R113 | F114 | K115 | I117 | T120 | Y121 | R123 | N154 |
| Rab5C | Q21 | K23 | V42 | K43 | E51 | I54 | G55 | A56 | F58 | L59 | T60 | E73 | W75 | E81 | R82 | Y83 | S85 | L86 | M89 | Y90 | R92 | A123 |
| Rab5B | Q20 | K22 | V41 | K42 | E50 | I53 | G54 | A55 | F57 | L58 | T59 | E72 | W74 | E80 | R81 | Y82 | S84 | L85 | M88 | Y89 | R91 | A122 |
| Rab9L | L7 | K9 | V28 | T29 | F37 | I40 | G41 | V42 | F44 | L45 | N46 | Q59 | W61 | E67 | R68 | F69 | S71 | L72 | P75 | Y77 | R78 | D110 |
| Rab6B | K13 | K15 | M34 | Y35 | Q43 | I46 | G47 | D49 | F50 | L51 | S52 | Q65 | W67 | E73 | R74 | F75 | S77 | L78 | S81 | Y82 | R84 | S117 |
| Rab18 | T7 | K10 | T29 | D31 | A38 | I41 | G42 | V43 | F45 | K46 | V47 | A60 | W62 | E68 | R69 | F70 | T72 | L73 | S76 | Y77 | R79 | T111 |
| Rab21 | S19 | F20 | C40 | E41 | I49 | L52 | Q53 | A54 | F56 | L57 | T58 | A71 | W73 | E79 | R80 | F81 | A83 | L84 | I87 | Y88 | R90 | L121 |
| Rab40B | | | | | | | | | | | | | | | | | | | | | | |
| Rab8B | D6 | K10 | S29 | E30 | I38 | I41 | G42 | D44 | F45 | K46 | I47 | Q60 | W62 | E68 | R69 | F70 | T72 | I73 | A76 | Y77 | R79 | A110 |
| Rab10 | L8 | K11 | S30 | D31 | I39 | I42 | G43 | I44 | F46 | K47 | I48 | Q61 | W63 | E69 | R69 | F71 | T73 | I74 | S77 | Y78 | R80 | A111 |
| Rab26 | V62 | K65 | K84 | D85 | I94 | V97 | G98 | I99 | F101 | R102 | N103 | Q116 | W118 | E124 | R125 | F126 | S128 | V129 | A132 | Y133 | R135 | A166 |
| Rab8A | L8 | K10 | S29 | D31 | I38 | I41 | G42 | D44 | F45 | K46 | I47 | Q60 | W62 | E68 | R69 | F70 | T72 | I73 | A76 | Y77 | R79 | A110 |
| Rab3A | M22 | K24 | A43 | D44 | V52 | V55 | G56 | D58 | F59 | K60 | V61 | Q74 | W76 | E82 | R83 | Y84 | T86 | I87 | A90 | Y91 | R93 | S124 |
| Rab27B | L9 | K11 | T30 | D31 | I39 | V42 | G43 | I44 | F46 | R47 | E48 | Q71 | W73 | E79 | R80 | F81 | S83 | L74 | A87 | F88 | R90 | A121 |
| Rab35 | L8 | K10 | A29 | D30 | I38 | I41 | G42 | V43 | F45 | K46 | R48 | Q60 | W62 | E68 | R69 | F70 | T72 | I73 | T76 | Y77 | R79 | C110 |
| Rab27A | L9 | K11 | T30 | D31 | I39 | V42 | G43 | I44 | F46 | R47 | E48 | Q71 | W73 | E79 | R80 | F81 | S83 | L84 | A87 | Y88 | R90 | A121 |
| Rab28 | Q12 | K14 | A33 | E35 | K42 | I45 | G46 | D48 | F50 | L51 | R52 | Q65 | W67 | G71 | Q72 | T73 | G75 | M77 | K81 | Y82 | Y84 | S118 |
| Rab33A | I36 | K38 | C57 | G58 | E66 | I69 | G70 | V71 | F73 | R74 | E75 | Q88 | W90 | E96 | R97 | F98 | K100 | M102 | H105 | Y106 | R108 | V140 |
| Rab30 | L9 | K11 | T30 | Q31 | E38 | I42 | G43 | V44 | F46 | M47 | I48 | Q61 | W63 | E69 | R70 | F71 | S73 | I74 | S77 | Y78 | R80 | S112 |
| Rab38 | E7 | L9 | V30 | H31 | A40 | I42 | G43 | V44 | F46 | A47 | L48 | L61 | W64 | E70 | R71 | F72 | N74 | M75 | V78 | Y79 | R81 | L114 |
| Rab24 | V6 | K9 | V28 | H29 | Q38 | I41 | G42 | A43 | F45 | V46 | A47 | G60 | W62 | E68 | R69 | Y70 | A72 | M73 | I76 | Y77 | R79 | E110 |
| Rab7L1 | D5 | L7 | S28 | Q29 | K37 | V40 | G41 | V42 | F44 | A45 | L46 | Q60 | W62 | E68 | R69 | F70 | S72 | L76 | Y77 | Y78 | R79 | L112 |
| Rab7A | L8 | K10 | V29 | N30 | K38 | I41 | G42 | A43 | F45 | L46 | T47 | Q60 | W62 | E68 | R69 | F70 | S72 | L73 | A76 | Y78 | R79 | A110 |
| Rab22B | I3 | L6 | V26 | Q27 | S35 | I38 | G39 | A40 | F42 | K45 | T46 | L57 | W59 | E65 | R66 | F67 | S69 | L70 | M73 | Y74 | R76 | P108 |
| Rab32 | E23 | L25 | V46 | H47 | R55 | I58 | G59 | F62 | L64 | K65 | V66 | L77 | L79 | E86 | R87 | M91 | R93 | V94 | K97 | E98 | A99 | L130 |
| Rab9A | L7 | K9 | V28 | T29 | F37 | I40 | G41 | V42 | F44 | L45 | N46 | Q59 | W61 | E67 | R68 | F69 | S71 | L72 | P75 | Y77 | R78 | D110 |
| RabL2B | | | | | | | | | | | | | | | | | | | | | | |
| Rab40C | | | | | | | | | | | | | | | | | | | | | | |
| Rab2A | | | | | | | | | | | | | | | | | | | | | | |
| Rab6C | | | | | | | | | | | | | | | | | | | | | | |
| Rab13 | | | | | | | | | | | | | | | | | | | | | | |
| Rab20 | | | | | | | | | | | | | | | | | | | | | | |

1

Supplementary Figure 10. Residues in ARF3 important for binding to autoprocessed MCF are functionally conserved in degraded, cleaved, and unaffected Rabs. Amino acid residues of ARF3 important for interacting with aMCF$^{CS}$ at top [1]. Residues that spatially aligned at these positions on the screened Rabs, in predicted Rab-aMCF complexes when overlayed onto ARF3-aMCF$^{CS}$ (PDB code 6ii6) [1], with these ARF3 residues were manually identified. Rabs degraded (names highlighted in red), cleaved (names in red font), and unaffected (names in black font) are denoted. For Rabs with no residues in the table, the Rab in the predicted co-structure with aMCF did not align in the same orientation as ARF3. Acidic (red), basic (blue), hydrophobic (gray), and hydrophilic (purple) amino acids are colored according to functionality.

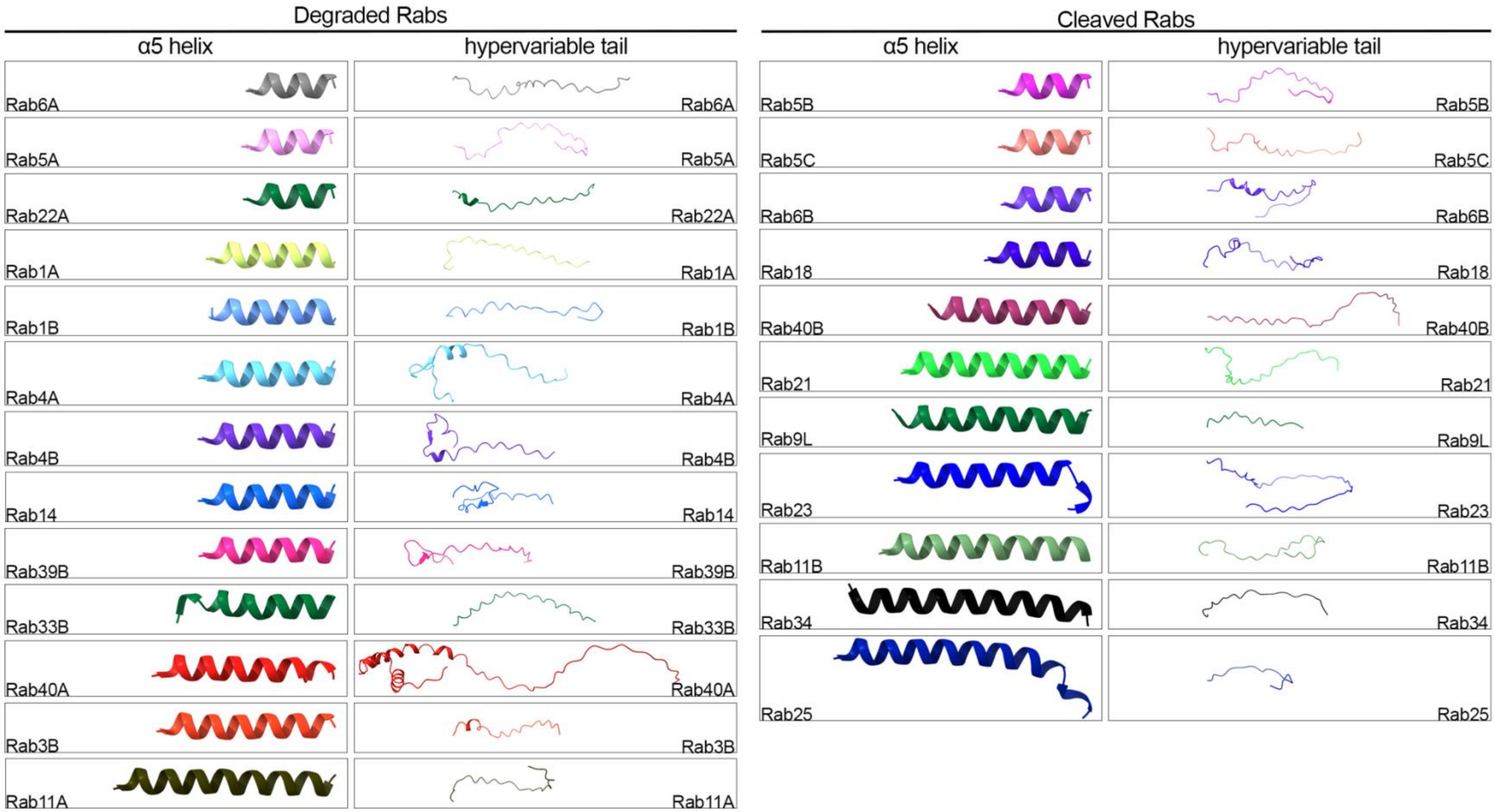
Degraded Rabs
α5 helix
hypervariable tail
Rab6A
Rab5A
Rab22A
Rab1A
Rab1B
Rab4A
Rab4B
Rab14
Rab39B
Rab33B
Rab40A
Rab3B
Rab11A
Cleaved Rabs
α5 helix
hypervariable tail
Rab5B
Rab5C
Rab6B
Rab18
Rab40B
Rab21
Rab9L
Rab23
Rab11B
Rab34
Rab25

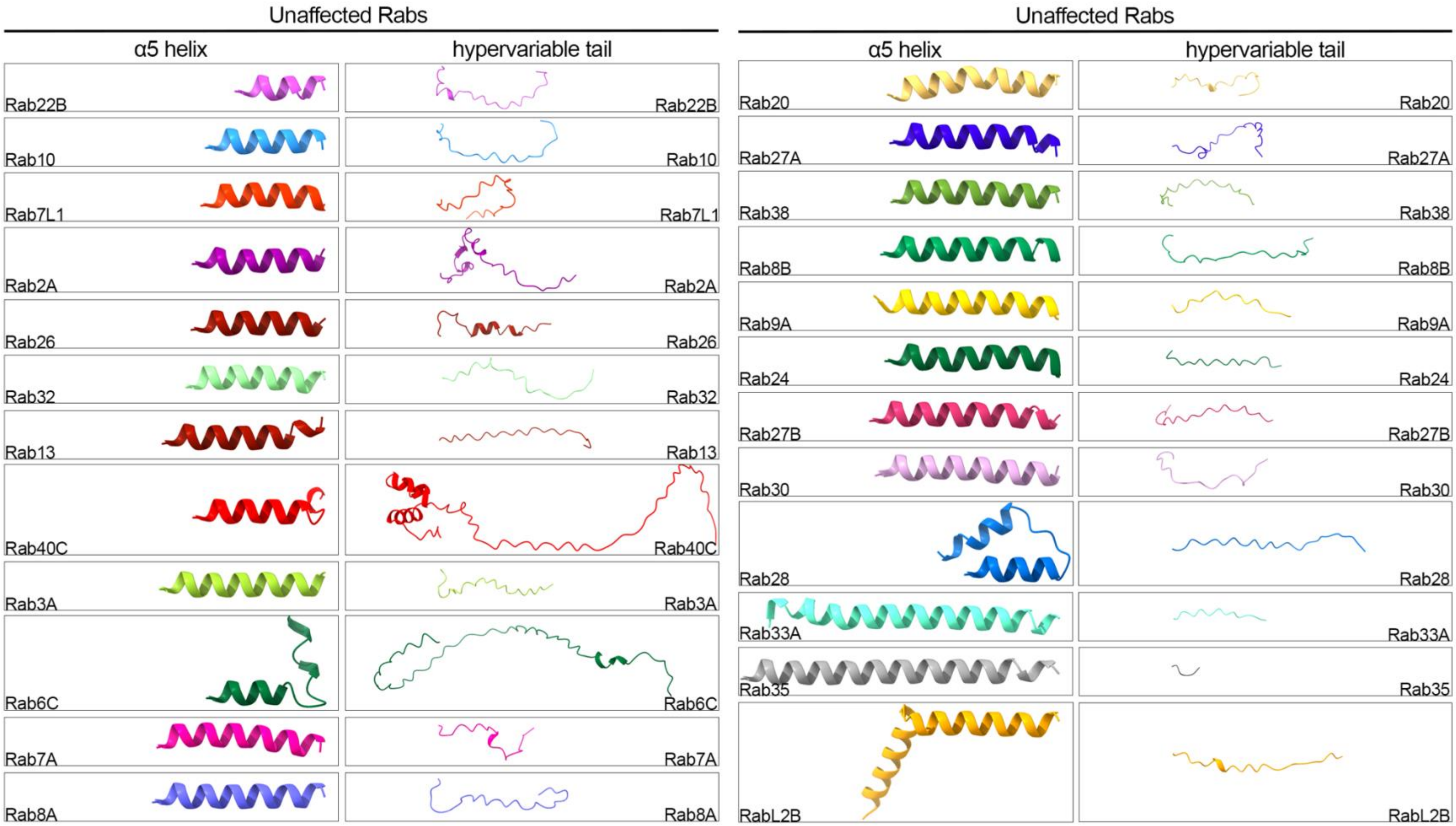


Supplementary Figure 11. Predicted structure at the fifth $\alpha$-helix and the hypervariable tail of Rabs. The fifth $\alpha$-helix ($\alpha$5) and hypervariable tail of Rabs used in the screen (Fig. 1b) was manually derived from the predicted co-structure of each Rab isoform with aMCF. Rabs categorized by those degraded, cleaved, and unaffected when co-expressed with MCF in order from shortest to longest $\alpha5$ within each category.

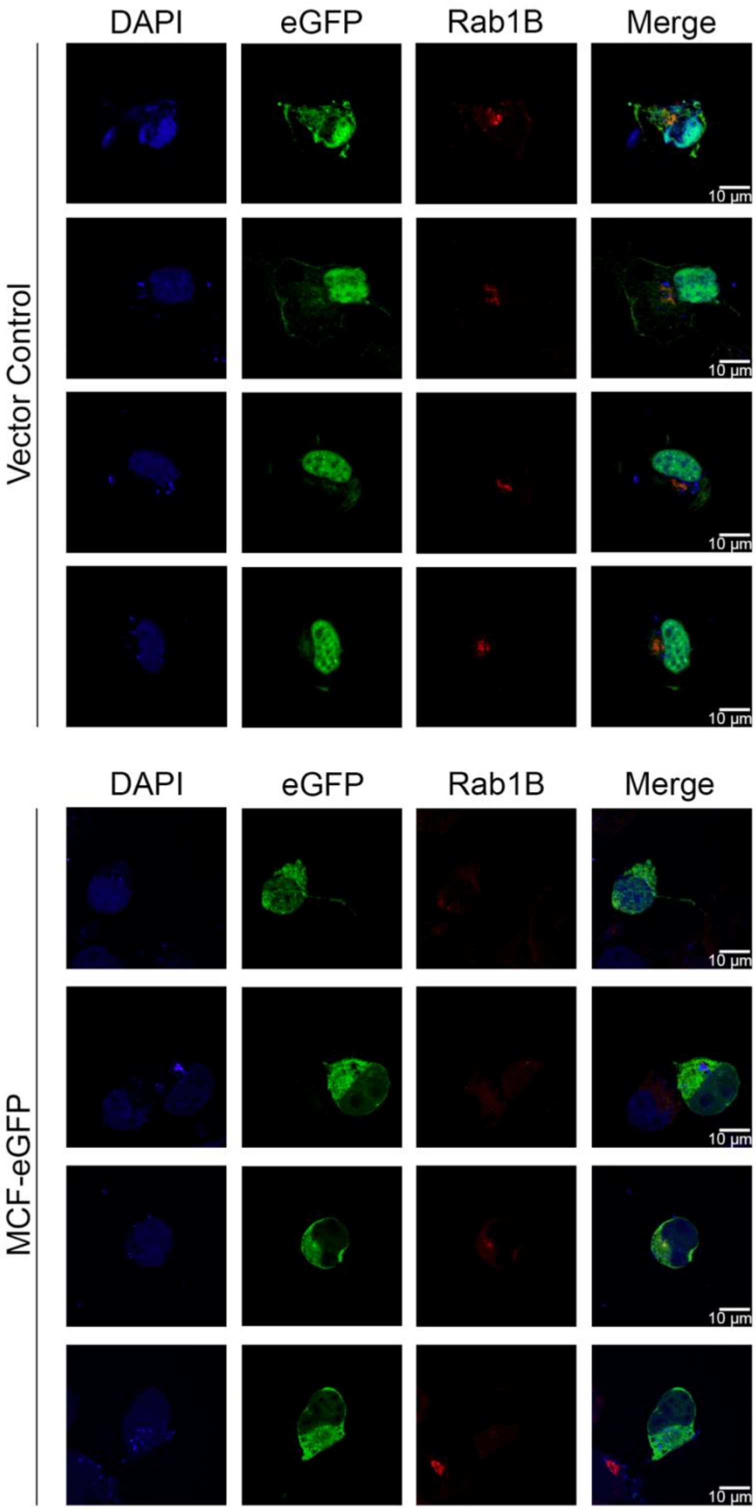


Supplementary Figure 12. Ectopic expression of MCF causes reorganization of endogenous Rab1B in Cos7 cells. Immunofluorescence microscopy shows endogenous Rab1B (red) when transfected with an empty GFP expression vector control (peGFP-N3) (top panel) or MCF-eGFP (bottom panel) (green). Nuclei is detected with 4′,6′-diamidino-2-phenylindole (DAPI) (in blue), Rab1B with anti-Rab1B antibodies, and MCF with anti-GFP antibodies.

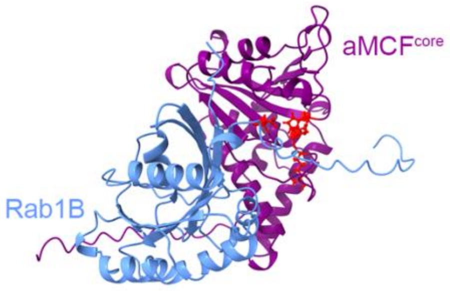


Supplementary Figure 13. Ribbon structure of predicted complex of aMCF$^{119-361}$ (purple) with Rab1B (cornflower blue). Residues important for the catalytic activity of aMCF shown in red.

Supplementary Table 1. Table of fluorescently-tagged Rab constructs used in co-expression screen with MCF.

| Rab | Vector background | Reference Information | Fifth Alpha Helix Sequence | Fifth Alpha Helix Length | Fifth Alpha Helix Angle | Tail Sequence | Tail Length |
|---|---|---|---|---|---|---|---|
| Rab1A | peGFP-C2 | Rzomp et al. 2003 | VEQSFMTMAAEIKKRM | 16 | 1.2 | GPGATAGGAEKSNVKIQSTPVKQSGGGCC | 29 |
| Rab4A | peGFP-C2 | Rzomp et al. 2003 | VEEAFVQCARKILNKIE | 17 | 7.3 | SGELDPERMGSGIQYGDAALRQLRSPRRAQAPNAQECGC | 39 |
| Rab4B | peGFP-C2 | Rzomp et al. 2003 | VEEAFLKCARTILNKID | 17 | 6.2 | SGELDPERMGSGIQYGDASLRQLRQPRSAQAVAPQPCGC | 39 |
| Rab6A | peGFP-C2 | Rzomp et al. 2003 | VKQLFRRVAAA | 10 | 1.8 | LPGMESTQDRSREDMIDIKLEKPQEQPVSEGGCSC | 35 |
| Rab11A | peGFP-C2 | Rzomp *et al.* 2003 | VEAAFQTILTEIYRIVSQKQMSDRREND | 28 | 3.7 | MSPSNNVVPIHVPPTTENKPKVQCCQNI | 28 |
| Rab14 | peGFP-C2 | provided by James Casanova | VEDAFLEAAKKIYQNIQ | 17 | 6.7 | DGSLDLNAAESGVQHKPSAPQGGRLTSEPQPQREGCGC | 38 |
| Rab22A | pEGFP-C3 | provided by Julie G. Donaldson at NIH/NHLBI | INELFIEISRR | 11 | 0.7 | IPSTDANLPSGGKGFKLRRQPSEPKRSCC | 29 |
| Rab3B | peCFP-C1 | Heo and Meyer 2003 | VRQAFERLVDAICDKMSDSLDT | 22 | 1.2 | DPSMLGSSKNTRLSDTPPLLQQNCSC | 26 |
| Rab1B | peGFP-C1 | Alvarez *et al.* 2003 | VEQAFMTMAAEIKKRM | 16 | 4.8 | GPGAASGGERPNLKIDSTPVKPAGGGCC | 28 |
| Rab33B | pEGFP-N1 | Smith *et al.* 2007 | PNDNDHVEAIFMTLAHKLKS | 20 | 9 | HKPLMLSQPPDNGIILKPEPKPAMTCWC | 28 |
| Rab39B | peGFP-C2 | Rzomp *et al.* 2003 | VEKAFTDLTRDIYELVK | 17 | 6.2 | RGEITIQEGWEGVKSGFVPNVVHSSEEVVKSERRCLC | 37 |
| *Rab40A* | pEGFP-N1 | Smith *et al.* 2007 | IIESFTELARIVLLRHRMNWLG | 21 | 0.9 | RPSKVLSLQDLCCRTIVSCTPVHLVDKLPLPSTLRSHLKSFSMAK GLNARMMRGLSYSLTTSSTHKSSLCKVEIVCPPQSPPKNCTRNS CKIS | 93 |
| Rab5A | peCFP-C1 | Heo *et al.* 2006 | VNEIFMAIAKK | 11 | 2.1 | LPKNEPQNPGANSARGRGVDLTEPTQPTRNQCCSN | 35 |
| Rab6B | peGFP-C2 | Rzomp et al. 2003 | VKQLFRRVASA | 11 | 2.8 | LPGMENVQEKSKEGMIDIKLDKPQEPPASEGGCSC | 35 |
| Rab5B | peCFP-C1 | Heo and Meyer 2003 | VNDLFLAIAKK | 11 | 1.7 | LPKSEPQNLGGAAGRSRGVDLHEQSQQNKSQCCSN | 35 |
| Rab34 | peGFP-C1 | Sun *et al.* 2003 | VREFFFRVAALTFEANVLAELEKSGARRIGD | 31 | 9.9 | VVRINSDDSNLYLTASKKKPTCCP | 24 |
| Rab25 | peGFP-C2 | provided by James Casanova | VELAFETVLKEIFAKVSKQRQNSIRTNAITLGS | 33 | 13.5 | AQAGQEPGPGEKRACCISL | 19 |
| Rab5C | peCFP-C1 | Heo and Meyer 2003 | VNEIFMAIAKK | 11 | 1.7 | LPKNEPQNATGAPGRNRGVDLQENNPASRSQCCSN | 35 |
| Rab9L | peCFP-C1 | Heo and Meyer 2003 | VTVAFEEAVRQVLAVEEQLEHCMLG | 25 | 0.3 | HTIDLNSGSKAGSSCC | 16 |
| Rab11B | peCFP-C1 | Heo and Meyer 2003 | VEEAFKNILTEIYRIVSQKQIADRAAH | 27 | 0.3 | DESPGNNVVDISVPPTTDGQKPNKLQCCQNL | 31 |
| Rab18 | peCFP-C1 | Heo and Meyer 2003 | VQCAFEELVEKII | 13 | 5.1 | QTPGLWESENQNKGVKLSHREEGQGGGACGGYCSVL | 36 |
| Rab21 | peCFP-C1 | Heo and Meyer 2003 | IEELFLDLCKRMIETAQVDERAKG | 24 | 4.5 | NGSSQPGTARRGVQIIDDEPQAQTSGGGCCSSG | 33 |
| Rab23 | peCFP-C1 | Heo and Meyer 2003 | VNEVFKYLAEKYLQKLKQQIAEPEL | 25 | 9.4 | THSSSNKIGVFNTSGGSHSGQNSGTLNGGDVINLRPNKQRTKKN RNPFSSCSIP | 53 |
| Rab40B | peCFP-C2 | Heo *et al.* 2006 | IALRSHLKSFSMANGLNARMM | 21 | 0.4 | HGGSYSLTTSSTHKRSSLRKVKLVRPPQSPPKNCTRNSCKIS | 42 |
| Rab3A | peCFP-C1 | Heo and Meyer 2003 | VKQTFERLVDVICEKMSESLD | 21 | 5.8 | TADPAVTGAKQGPQLSDQQVPPHQDCAC | 28 |
| Rab27A | peCFP-C1 | Heo and Meyer 2003 | ISQAIEMLLDLIMKRMERCVDK | 22 | 4.8 | SWIPEGVVRSNGHASTDQLSEEKEKGACGC | 30 |
| Rab7A | pEGFP-C3 | Feng *et al.* 2001 | VEQAFQTIARNALKQETEVELY | 22 | 1.3 | NEFPEPIKLDKNDRAKASAESCSC | 24 |
| Rab9A | pEGFP-C3 | Barbero *et al.* 2002 | VAAAFEEAVRRVLATEDRSDHLI | 23 | 7.5 | QTDTVNLHRKPKPSSSCC | 18 |
| Rab10 | peGFP-C2 | Rzomp et al. 2003 | IEKAFLTLAEDILRK | 15 | 0.5 | TPVKEPNSENVDISSGGGVTGWKSKCC | 27 |
| Rab2A | peCFP-C1 | Heo and Meyer 2003 | VEEAFINTAKEIYEKIQ | 17 | 6.5 | EGVFDINNEANGIKIGPQHAATNATHAGNQGGQQAGGGCC | 40 |
| Rab6C | peCFP-C1 | Heo and Meyer 2003 | VKQLFRRVAAALPGMESTQDGS | 22 | 44.4 | REDMSDIKLEKPQEQTVSEGGCSCYSPMSSSTLPQKPPYSFIDC SVNIGLNLFPSLITFCNSSLLPVSWR | 69 |
| Rab7L1 | peCFP-C1 | Heo and Meyer 2003 | INEAMRVLIEKMMRNS | 16 | 0.5 | TEDIMSLSTQGDYINLQTKSSSWSCC | 26 |
| Rab8A | peCFP-C1 | Heo and Meyer 2003 | VENAFFTLARDIKAKMDKKLEG | 22 | 1.8 | NSPQGSNQGVKITPDQQKRSSFFRCVLL | 28 |
| Rab8B | peCFP-C1 | Heo and Meyer 2003 | VEEAFFTLARDIMTKLNRKMNDS | 23 | 0.5 | NSAGAGGPVKITENRSKKTSFFRCSLL | 27 |
| Rab13 | peCFP-C1 | Heo and Meyer 2003 | VDEAFSSLARDILLKSGGRR | 20 | 9.6 | SGNKPPSTDLKTCDKKNTNKCSLG | 24 |
| Rab22B | peCFP-C1 | Heo and Meyer 2003 | IEELFQGISRQ | 11 | 0.7 | IPPLDPHENGNNGTIKVEKPTMQASRRCC | 29 |
| Rab26 | peCFP-C1 | Heo and Meyer 2003 | VDLAFTAIAKELKQRSM | 17 | 4.8 | KAPSEPRFRLHDYVKREGRGASCCRP | 26 |

| | | | | | | | |
|---|---|---|---|---|---|---|---|
| Rab27B | peCFP-C1 | Heo and Meyer 2003 | VEKAVETLLDLIMKRMEQCVEKTQ | 24 | 1.6 | IPDTVNGGNSGNLDGEKPPEKKCIC | 25 |
| Rab28 | peCFP-C1 | Heo and Meyer 2003 | VFLCFQKVAAEILGIKLNKAEIEQSQRIV | 29 | 41.9 | RAEIVKYPEEENQHTTSTQSRICSVQ | 25 |
| Rab30 | peCFP-C1 | Heo and Meyer 2003 | VEKLFLDLACRLISEARQNTLVNN | 24 | 3 | VSSPLPGEGKSISYLTCCNFN | 21 |
| Rab32 | peCFP-C1 | Heo and Meyer 2003 | IEEAARFLVEKILVNHQS | 18 | 3.1 | FPNEENDVDKIKLDQETLRAENKSQCC | 27 |
| Rab33A | peCFP-C1 | Heo and Meyer 2003 | PKESQNVESIFMCLACRLKAQKSLLYRDAERQQGKVQK | 38 | 1.8 | LEFPQEANSKTSCPC | 15 |
| Rab35 | peCFP-C1 | Heo and Meyer 2003 | VEEMFNCITELVLRAKKDNLAKQQQQQQNDVVKLTKNSKR | 40 | 1 | KKRCC | 5 |
| Rab38 | peCFP-C1 | Heo and Meyer 2003 | IDEASRCLVKHILANECDLMES | 22 | 0.2 | IEPDVVKPHLTSTKVASCSGCAKS | 24 |
| RabL2B | peCFP-C1 | Heo and Meyer 2003 | VVKLFNDAIRLAVSYKQNSQDFMDEIFQELENFSLEQEEE | 40 | 26.6 | DVPDQEQSSSIETPSEEAASPHS | 23 |
| Rab40C | peGFP-N1 | Smith *et al.* 2007 | VIESFTELSRIVLMRHGMEK | 20 | 8.2 | IWRPNRVFSLQDLCCRAIVSCTPVHLIDKLPLPVTIKSHLKSFSMA NGMNAVMMHGRSYSLASGAGGGGSKGNSLKRSKSIRPPQSPP QNCSRSNCKIS | 99 |
| Rab20 | peGFP-N1 | Smith *et al.* 2007 | VDLLFETLFDLVVPMILQQRAE | 22 | 3.3 | RPSHTVDISSHKPPKRTRSGCCA | 23 |
| Rab24 | peGFP-C1 | Munafó and Colombo 2002 | VDELFQKVAEDYVSVAAFQVMTE | 23 | 5 | DKGVDLGQKPNPYFYSCCHH | 20 |

Rabs unaffected (names in black font), cleaved (names in red font), and degraded (names highlighted in red) when co-expressed with MCF in HEK 293T cells categorized. Table includes the vector background each Rab isoform is expressed in, as well as the length and amino acid sequence of their fifth α-helix and C-terminal tail. Angle of α5 helix of each Rab measured as the angle from the N- to the C-terminus of the helix. The reference information for each construct is also include

| Supplementary Table 2. Sequences of gBlocks used for plasmid construction. | | |
|---|---|---|
| gBlock | Sequence | Vector |
| aMCFdomain II (residues 85-324) | CCTGTACTTCCAATCCAATGCTATGGTGACGTTCCAGAACAAGTCTGAGAAGTACAACCGATTGTTCCG<br>TGAGATTGCTTCTGCTGGCGTGGTGGATGCGAAAGCGACTGAACAGCTTGCGCCACAGTTAATGCTG<br>CTGAACCTATCGAATGACGGTTTTGGTGGGCGTTGTGATCCACTTTCTAAACTCGTTTTGGTTGCGAAA<br>CAGCTTGAAAACGATGGTCAAGTTGGCGTGGCAAGACAACTGCTAGAAAAGATGTACTCTGCGGCAGC<br>GGTGCTGAGCAATCCAACCCTTTACTCAGACAGTGAAAAAGCCAATGCAAGCAAGTTGCTCAGCAGCT<br>TGGCGGCCATTCATGCGAAGAACCCAATGCATGATACGTCGATGAAAGTGTGGCAGGAAAAGCTGGA<br>AGGGAAGCAAGCGCTGACCGTAAACGGTGTGGTTGAGAAAATCACTGATGCATCGGCTAACGGTAAA<br>CCTGTGCTGTTGGAACTTGATGCTCCGGGGCATGCGATGGCAGCTTGGGCAAAAGGCTCAGGCGACG<br>ATCGTGTTTACGGCTTCTACGATCCAAATGCTGGCATCGTTGAGTTTTCGTCAGCAGAGAAGTTTGGC<br>GACTACCTAACGCGTTTCTTCGGCAAGTCCGATCTGAACATGGCTCAAAGCTATAAGCTGGGTAAAAA<br>CGACGCAGGTGAAGCAATCTTCAACCGCGTGGTGGTAATGGATGGCAATACATTAGCAAGCTACAAGT<br>AAATTGGAAGTGGATAACGG | pMCSG7 |
| aMCFdomain I (residues 1-84) | CCTGTACTTCCAATCCAATGCTATGGGACTAGAGAAAGACTTTAAACGCTATGGCGACGCGCTGAAAC<br>CAGATACGAGCGTGCCGGGTAAATCGAAAGACATTCGCACCACTAAAGATTTCCTAAATGGTTACAAAA<br>ATGACCATGCGAAAGAGATCGTTGACGGCTTCCGCTCAGATATGAGTATCAAGCAACTGGTGGATCTG<br>TTTGTTAAAGGTAACTGGAGTGCAGAGCAAAAAGGTGCGCTTGCTTGGGAAATCGAAAGTCGTGCACT<br>GAAAGAACAAAAACTCATCTCAGAAGAGGATCTGTAAATTGGAAGTGGATAACGG | pMCSG7 |
| Rab1B | TCCGGTGGTGGTGGTGGAATTATGAACCCCGAATATGACTACCTGTTTAAGCTGCTTTTGATTGGCGA<br>CTCAGGCGTGGGCAAGTCATGCCTGCTCCTGCGGTTTGCTGATGACACGTACACAGAGAGCTACATC<br>AGCACCATCGGGGTGGACTTCAAGATCCGAACCATCGAGCTGGATGGCAAAACTATCAAACTTCAGAT<br>CTGGGACACAGCGGGCCAGGAACGGTTCCGGACCATCACTTCCAGCTACTACCGGGGGGCTCATGG<br>CATCATCGTGGTGTATGACGTCACTGACCAGGAATCCTACGCCAACGTGAAGCAGTGGCTGCAGGAG<br>ATTGACCGCTATGCCAGCGAGAACGTCAATAAGCTCCTGGTGGGCAACAAGAGCGACCTCACCACCA<br>AGAAGGTGGTGGACAACACCACAGCCAAGGAGTTTGCAGACTCTCTGGGCATCCCCTTCTTGGAGAC<br>GAGCGCCAAGAATGCCACCAATGTCGAGCAGGCGTTCATGACCATGGCTGCTGAAATCAAAAAGCGG<br>ATGGGGCCTGGAGCAGCCTCTGGGGGCGAGCGGCCCAATCTCAAGATCGACAGCACCCCTGTAAAG<br>CCGGCTGGCGGTGGCTGTTGCTAGCTAGACTCCATGGGTCGACTCGAGC | pGEX-KG |
| Rab23 | TCCGGTGGTGGTGGTGGAATTTCTAGAGAAAATTTATATTTTCAAGGTATGTTGGAGGAAGATATGGAA<br>GTCGCCATAAAGATGGTGGTTGTAGGGAATGGAGCAGTTGGAAAATCAAGTATGATTCAGCGATATTG<br>CAAAGGCATTTTTACAAAAGACTACAAGAAAACCATTGGAGTTGATTTTTTGGAGCGACAAATTCAAGTT<br>AATGATGAAGATGTCAGACTAATGTTATGGGACACTGCAGGTCAGGAGGAATTTGATGCAATTACAAAG<br>GCCTACTATCGAGGAGCCCAGGCTTGTGTGCTCGTGTTCTCTACCACAGATAGGGAATCTTTTGAAGC<br>AGTTTCCAGTTGGAGAGAGAAAGTAGTAGCCGAAGTGGGAGATATACCAACTGTACTTGTGCAAAACA<br>AGATTGATCTTCTGGATGATTCTTGTATAAAGAATGAGGAAGCTGAGGCACTGGCAAAAAGGTTAAAGT<br>TAAGATTCTACAGAACATCAGTGAAAGAAGATCTAAATGTGAATGAAGTTTTTAAGTATTTGGCTGAAAA<br>ATACCTTCAGAAACTCAAACAACAAATAGCTGAGGATCCAGAACTAACGCATTCAAGTAGTAACAAGAT<br>TGGTGTCTTTAATACATCTGGTGGAAGTCACTCCGGTCAGAATTCAGGTACCCTCAATGGTGGAGATG<br>TCATCAATCTTAGACCCAACAAACAAAGGACCAAGAAAAACAGAAATCCTTTTAGCAGCTGTAGCATAC<br>CCTAGTCTAGACTAGACTCCATGGGTCGACTCGAGC | pGEX-KG |
| Rab2A | CATGGACGAGCTGTACAAGTCCGGCCGGACTCAGATCATGGCGTACGCCTATCTCTTCAAGTACATCA<br>TAATCGGCGACACAGGTGTTGGTAAATCATGCTTATTGCTACAGTTTACAGACAAGAGGTTTCAGCCAG<br>TGCATGACCTTACTATTGGTGTAGAGTTCGGTGCTCGAATGATAACTATTGATGGGAAACAGATAAAAC<br>TTCAGATATGGGATACGGCAGGGCAAGAATCCTTTCGTTCCATCACAAGGTCGTATTACAGAGGTGCA<br>GCAGGAGCTTTACTAGTTTACGATATTACACGGAGAGATACATTCAACCACTTGACAACCTGGTTAGAA<br>GATGCCCGCCAGCATTCCAATTCCAACATGGTCATTATGCTTATTGGAAATAAAAGTGATTTAGAATCTA<br>GAAGAGAAGTAAAAAAAGAAGAAGGTGAAGCTTTTGCACGAGAACATGGACTCATCTTCATGGAAACG<br>TCTGCTAAGACTGCTTCCAATGTAGAAGAGGCATTTATTAATACAGCAAAAGAAATTTATGAAAAAATTC<br>AAGAAGGAGTCTTTGACATTAATAATGAGGCAAATGGCATTAAAATTGGCCCTCAGCATGCTGCTACCA<br>ATGCAACACATGCAGGCAATCAGGGAGGACAGCAGGCTGGGGGCGGCTGCTGTTGATCGAGCTCAA<br>GCTTCGAATTCTGCAGTCGACGGTA | peGFP-C2 |
| Rab6C | CATGGACGAGCTGTACAAGTCCGGCCGGACTCAGATCATGTCCGCGGGCGGAGACTTCGGGAATCC<br>GCTGAGGAAATTCAAGCTGGTGTTCCTGGGGGAGCAAAGCGTTGCAAAGACATCTTTGATCACCAGAT<br>TCAGGTATGACAGTTTTGACAACACCTATCAGGCAATAATTGGCATTGACTTTTTATCAAAAACTATGTA<br>CTTGGAGGATGGAACAATCGGGCTTCGGCTGTGGGATACGGCGGGTCAGGAACGTCTCCGTAGCCTC<br>ATTCCCAGGTACATCCGTGATTCTGCTGCAGCTGTAGTAGTTTACGATATCACAAATGTTAACTCATTC<br>CAGCAAACTACAAAGTGGATTGATGATGTCAGAACAGAAAGAGGAAGTGATGTTATCATCACGCTAGTA<br>GGAAATAGAACAGATCTTGCTGACAAGAGGCAAGTGTCAGTTGAGGAGGGAGAGAGGAAAGCCAAAG<br>GGCTGAATGTTACGTTTATTGAAACTAGGGCAAAAGCTGGATACAATGTAAAGCAGCTCTTTCGACGTG<br>TAGCAGCAGCTTTGCCGGGAATGGAAAGCACACAGGACGGAAGCAGAGAAGACATGAGTGACATAAA<br>ACTGGAAAAGCCTCAGGAGCAAACAGTCAGCGAAGGGGGTTGTTCCTGCTACTCTCCCATGTCATCTT<br>CAACCCTTCCTCAGAAGCCCCCTTACTCTTTCATTGACTGCAGTGTGAATATTGGCTTGAACCTTTTCC<br>CTTCATTAATAACGTTTTGCAATTCATCATTGCTGCCTGTCTCGTGGAGGTGATCGAGCTCAAGCTTCG<br>AATTCTGCAGTCGACGGTA | peGFP-C2 |

| | | |
|---|---|---|
| Rab35 | CATGGACGAGCTGTACAAGTCCGGCCGGACTCAGATCATGGCCCGGGACTACGACCACCTCTTCAAG<br>CTGCTCATCATCGGCGACAGCGGTGTGGGCAAGAGCAGTTTACTGTTGCGTTTTGCAGACAACACTTT<br>CTCAGGCAGCTACATCACCACGATCGGAGTGGATTTCAAGATCCGGACCGTGGAGATCAACGGGGAG<br>AAGGTGAAGCTGCAGATCTGGGACACAGCGGGGCAGGAGCGCTTCCGCACCATCACCTCCACGTATT<br>ATCGGGGGACCCACGGGGTCATTGTGGTTTACGACGTCACCAGTGCCGAGTCCTTTGTCAACGTCAA<br>GCGGTGGCTTCACGAAATCAACCAGAACTGTGATGATGTGTGCCGAATATTAGTGGGTAATAAGAATG<br>ACGACCCTGAGCGGAAGGTGGTGGAGACGGAAGATGCCTACAAATTCGCCGGGCAGATGGGCATCC<br>AGTTGTTCGAGACCAGCGCCAAGGAGAATGTCAACGTGGAAGAGATGTTCAACTGCATCACGGAGCT<br>GGTCCTCCGAGCAAAGAAAGACAACCTGGCAAAACAGCAGCAGCAACAACAGAACGATGTGGTGAAG<br>CTCACGAAGAACAGTAAACGAAAGAAACGCTGCTGCTAATCGAGCTCAAGCTTCGAATTCTGCAGTCG<br>ACGGTA | peGFP-C2 |

Supplementary Table 3. X-ray data collection and refinement statistics for aMCF$^{CA}$ structure.

| **PDB Accession Code** | 8SFG |
|---|---|
| **Data Collection** | |
| Space group | $P2_1$ |
| Unit cell parameters (Å; º) | 61.38, 160.66, 71.79, 90.00, 90.00, 90.00 |
| Resolution range (Å) | 30.00 – 2.80 (2.85 – 2.80) |
| No. of reflections | 32763 (1579) |
| $R_{merge}$ (%) | 11.00 (75.10) |
| Completeness (%) | 97.7 (94.7) |
| $\langle I/\sigma(I)\rangle$ | 13.3 (1.9) |
| Multiplicity | 4.4 (4.3) |
| Wilson *B* factor | 54.5 |
| **Structure Determination** | |
| MR initial model (PDB code) | 6ii6 |
| **Refinement** | |
| Resolution range (Å) | 29.87 – 2.80 (2.87 – 2.80) |
| Completeness (%) | 97.5 (93.2) |
| No. of reflections | 31033 (2291) |
| $R_{work}/R_{free}$, (%) | 18.9/23.0 (26.2/29.6) |
| Protein chains/atoms | 4/10149 |
| Ligand/Solvent atoms | 128/83 |
| Mean temperature factor (Å$^2$) | 65.0 |
| **Coordinate Deviations** | |
| R.m.s.d. bonds (Å) | 0.004 |
| R.m.s.d. angles (º) | 1.203 |
| **Ramachandran plot** | |
| Favored (%) | 97.0 |
| Allowed (%) | 3.0 |
| Outside allowed (%) | 0.0 |
